# Clamp the LAMP: a photoelectrochemical platform for *KRAS* mutation detection via wild-type blocking

**DOI:** 10.64898/2026.02.22.707251

**Authors:** J. Strmiskova, A. Valverde, L. Moranova, J. Arnouts, F. Zavadil-Kokas, S. Koljenovic, K. Zwaenepoel, T. Vandamme, M. Bartosik, K. De Wael

**Affiliations:** Research Centre for Applied Molecular Oncology (RECAMO), Masaryk Memorial Cancer Institute, 65653 Brno, Czech Republic; National Centre for Biomolecular Research, Faculty of Science, Masaryk University, 62500 Brno, Czech Republic; Antwerp Engineering, Photoelectrochemistry and Sensing (A-PECS), University of Antwerp, 2020 Antwerp, Belgium; NANOlight Centre of Excellence, University of Antwerp, 2020 Antwerp, Belgium; Department of Pathology, Antwerp University Hospital (UZA), 2650 Edegem, Belgium; Department of Oncology, Antwerp University Hospital (UZA), 2650 Edegem, Belgium; Centre for Oncological Research (CORE), University of Antwerp, 2610 Wilrijk, Belgium

**Keywords:** *KRAS* mutation detection, clamp-inhibited LAMP, locked nucleic acid (LNA), wild-type suppression, singlet oxygen, photoelectrochemical biosensing, decentralized testing

## Abstract

*KRAS* mutations are among the most prevalent oncogenic alterations in colorectal, lung, and pancreatic cancer, yet their detection remains analytically challenging in the presence of an overwhelming wild-type (WT) background. Here, we report a photoelectrochemical (PEC) genotyping platform that integrates clamp-inhibited loop-mediated isothermal amplification (C-LAMP) with enzyme-free singlet oxygen (^1^O_2_)-driven PEC transduction for mutation-selective *KRAS* detection. Locked nucleic acid (LNA) clamp probes selectively suppress WT amplification during isothermal amplification, enriching mutant alleles and enabling single-nucleotide variant (SNV) discrimination with high selectivity. Amplified products are magnetically captured and transduced into photocurrent via visible-light-induced ^1^O_2_ redox cycling, eliminating enzymatic reporters and reducing background interference. The C-LAMP/PEC platform achieves a limit of detection of 35 copies µL^-1^ (58 aM) and a minimum detectable variant allele frequency (VAF) of 4.8% in heterogeneous mutant/WT genomic DNA mixtures. Analytical performance was validated in cancer cell lines and in patient-derived fresh frozen tissues, showing complete concordance with Nanopore sequencing and droplet digital PCR (ddPCR) within the evaluated cohort (n = 16). This work introduces a robust and modular PEC biosensing strategy that combines molecular WT suppession with enzyme-free photoelectrochemistry, offering an economically competitive and instrumentation-simplified approach for clinically relevant *KRAS* mutation analysis toward decentralized testing.

## 1. Introduction

Mutations in the *KRAS* (Kirsten rat sarcoma viral oncogene homolog) gene are among the most frequent oncogenic alterations in human cancers and represent key therapeutic targets (Yang et al. 2023). Together with *NRAS* and *HRAS*, *KRAS* encodes a small GTP-binding protein that regulates the Ras/Raf/mitogen-activated protein kinase (MAPK) pathway, a central mediator of cell proliferation and survival. Oncogenic *KRAS* mutations disrupt intrinsic GTPase activity, leading to persistent activation and tumorigenesis (Jancik et al. 2010). Such mutations occur in approximately 23% of adult cancers, predominantly as single-nucleotide variants (SNVs) at codons 12 and 13. Clinically relevant variants such as G12C and G12V are prevalent in colorectal cancer (CRC), non-squamous non-small cell lung cancer (NSCLC) and pancreatic ductal adenocarcinoma (PDAC), where they correlate with poor prognosis and reduced therapeutic response (Lee et al. 2022a; Meng et al. 2021; Tan and Tan 2022). Beyond prognostic significance, *KRAS* mutation status dictates eligibility for targeted therapies, including EGFR inhibitors and, more recently, pan-*KRAS* and allele-specific inhibitors currently advancing in clinical trials (Biller and Schrag 2021; Fu et al. 2021; Janes et al. 2018; Negri et al. 2022; Strickler et al. 2023; Zhao et al. 2017).

These considerations highlight the need for accurate and accessible *KRAS* genotyping technologies. Polymerase chain reaction (PCR) and next-generation sequencing (NGS) remain clinical standards for mutation detection, however, their widespread implementation is limited by instrumentation requirements, multi-step workflows and dependence on centralized facilities and trained personnel (Arnouts et al. 2025; Gilson et al. 2019; Sherwood et al. 2017). Modified PCR-based strategies, including peptide nucleic acid (PNA) blockers, co-amplification at lower denaturation temperature (COLD-PCR) and multiplex fluorescent assays, have improved allele enrichment and analytical sensitivity, but remain operationally complex (Carotenuto et al. 2012; Laosinchai-Wolf et al. 2011; Oh et al. 2010). Consequently, isothermal amplification techniques (IATs), which operate at constant temperature and reduce instrumentation demands, have gained increasing attention. Methods such as loop-mediated isothermal amplification (LAMP), rolling circle amplification (RCA) and recombinase polymerase amplification (RPA) have been explored for the detection of *KRAS* or other SNVs, frequently coupled with biosensing interfaces to simplify workflows and shorten assay times (He et al. 2025; Islam et al. 2025; Ji et al. 2025; Lazaro et al. 2022; Lee et al. 2022b; Li et al. 2022; Martorell et al. 2019; Mirlohi et al. 2024; Moranova et al. 2024; Ondraskova et al. 2023; Sebuyoya et al. 2025; Sebuyoya et al. 2023; Zhou et al. 2022). Among these, LAMP offers high amplification efficiency, robustness and compatibility with simplified hardware, making it attractive for decentralized molecular diagnostics.

Despite these advances, the detection of *KRAS* SNVs remains particularly challenging due to the overwhelming abundance of wild-type (WT) sequences, typically representing >99% of total genomic DNA in clinical samples (Franklin et al. 2010). This WT background severely compromises assay specificity at low variant allele frequencies (VAF), limiting reliable discrimination of rare mutant alleles and clinical translation of IAT-based biosensors. Locked nucleic acid (LNA)-modified probes have improved mismatch discrimination (Choate et al. 2024; Moranova et al. 2024), yet effective WT suppression under isothermal conditions remains difficult. In parallel, photoelectrochemical (PEC) biosensing has emerged as a powerful strategy for nucleic acid detection, combining high sensitivity, low background and compatibility with miniaturized instrumentation. Singlet oxygen (^1^O_2_)-driven PEC approaches leverage visible-light activation of photosensitizers to generate reactive oxygen species that trigger redox cycling of mediators such as hydroquinone (HQ), enabling enzyme-free photocurrent transduction (Trashin et al. 2017). While this concept offers improved stability and reduced reagent cost relative to enzymatic detection systems, previous ^1^O_2_-driven PEC studies have been primarily demonstrated using synthetic oligonucleotides or limited biological validation and have not been systematically applied to clinically relevant *KRAS* SNV genotyping (Daems et al. 2024; Shanmugam et al. 2024; Stratulat et al. 2025).

Here, we introduce a photoelectrochemical platform that integrates clamp-inhibited LAMP (C-LAMP) with ^1^O_2_-driven PEC detection for mutation-selective *KRAS* genotyping. The system employs LNA-modified clamp probes to block WT sequences during isothermal amplification, enriching mutant alleles and enabling single-nucleotide discrimination. The resulting amplificon are captured on magnetic microbeads functionalized with LNA-modified mutation-specific capture probes and transduced into stable photocurrents via visible-light-driven ^1^O_2_ redox chemistry. The architecture is inherently sequence-adaptable through probe redesign, providing an enzyme-free and flexible analytical framework for *KRAS* mutation detection with potential for decentralized implementation.

## 2. Results

### 2.1 Design and working principle of the C-LAMP/^1^O_2_-driven PEC platform

The proposed biosensing platform integrates C-LAMP with enzyme-free ^1^O_2_-driven PEC detection to enable SNV discrimination in the *KRAS* gene **(Figure 1)**. During amplification, two LNA-modified clamp probes (CL1 and CL2) hybridize to mutation hotspots at codons 12 and 13 of the *KRAS* WT sequence. The high duplex stability of LNA clamp probes selectively inhibits polymerase extension of perfectly matched WT templates during isothermal amplification, whereas single-nucleotide mismatches at mutant positions reduce clamp-target stability, permitting preferential amplification of mutant alleles. The resulting C-LAMP amplicons are captured by mutation-specific biotinylated LNA-modified capture probes (bCPs) immobilized on streptavidin-coated magnetic microbeads (Strep-MBs) and subsequently hybridized with detection probes (DPs) labeled with the photosensitizer chlorin e6 (Ce6). Upon 660 nm illumination, Ce6 generates ^1^O_2_ with a reported quantum yield of 0.71, promoting oxidation of the redox mediator hydroquinone (HQ) into benzoquinone (BQ) (Daems et al. 2024; Shanmugam et al. 2022; Stratulat et al. 2025). Although ^1^O_2_ is a reactive oxygen species, its generation is temporally confined to the final readout step (i.e., oxidation of HQ) and spatially restricted by its short lifetime and limited diffusion radius in aqueous media (Shanmugam et al. 2022), minimizing any impact on amplification or hybridization process.

**Figure 1.**
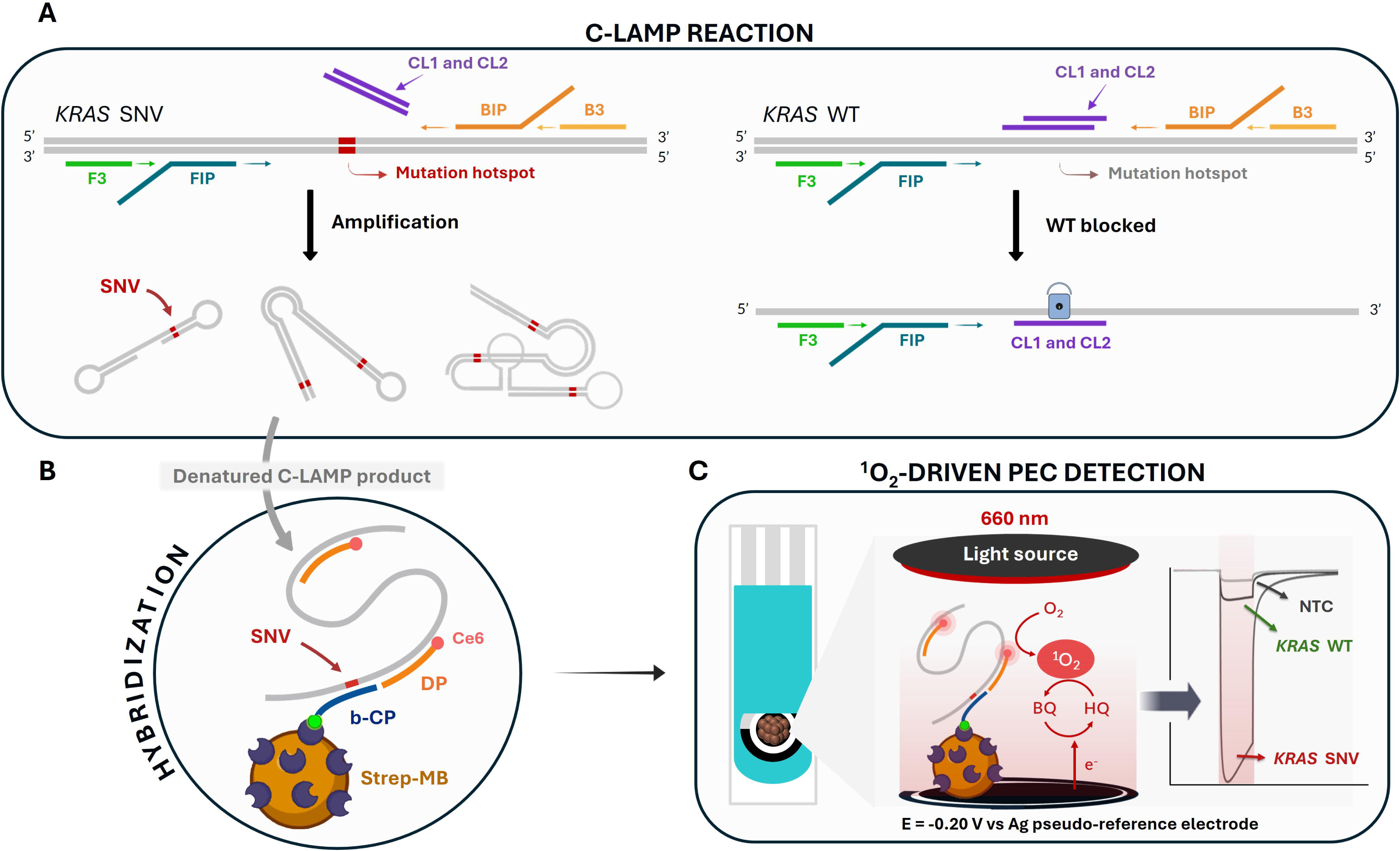
**(A)** Schematic illustration of the workflow integrating C-LAMP reaction with ^1^O_2_-driven PEC detection. LNA clamp probes hybridize to the *KRAS* WT sequence at codons 12-13 to suppress its amplification, while mutant templates are efficiently amplified. **(B)** Strep-MBs functionalized with b-CPs hybridize to the denatured C-LAMP amplicons together with Ce6-labeled DPs. **(C)** Under 660 nm illumination, Ce6 produces ^1^O_2_ that oxidizes HQ to BQ, generating photocurrent at the working surface of a screen-printed carbon electrode for unambiguous *KRAS* SNV/WT discrimination. NTC = non-template control.

Under an applied potential (−0.20 V versus Ag pseudo-reference electrode), HQ/BQ redox cycling produces a stable photocurrent at the screen-printed carbon electrode, providing a quantitative PEC signal proportional to the amount of mutation-specific amplicons. By combining molecular-level WT blocking during amplification with enzyme-free ^1^O_2_-driven PEC transduction, this integrated platform is designed to enhance analytical selectivity and robustness for *KRAS* mutation detection.

### 2.2 Optimization of C-LAMP and ^1^O_2_-driven PEC parameters

To establish optimal conditions for ^1^O_2_-driven PEC detection, synthetic *KRAS* G12V oligonucleotides were initially employed to compare magnetic bead scaffolds **(Figure S1)**. Strep-MBs functionalized with b-CPs were evaluated against carboxylated beads carrying amino-modified capture probes (HOOC-MB/a-CPs) (Daems et al. 2024; Moranova et al. 2024). Although both configurations generated measurable photocurrents in the presence of synthetic G12V targets, the Strep-MB/b-CPs platform provided higher signal-to-blank ratios and a lower limit of detection (39 pM vs 98 pM, blank defined as target-free control). This configuration was therefore selected for all subsequent experiments.

C-LAMP conditions were then optimized using genomic DNA (gDNA) extracted from cancer cell lines with *KRAS* genotypes confirmed by Nanopore sequencing **(Figure S2)**. As shown in **Table S1**, homozygous mutant (SW620, G12V/G12V), heterozygous mutant (SW837, G12C/WT), WT (HT-29, A2780) and non-template controls (NTC) were selected to rigorously maximize WT suppression while preserving SNV amplification across different mutant allele frequencies and cell lines. Optimal conditions consisted of 0.6 µM of each clamp probe (CL1 and CL2), an amplification temperature of 62 °C and a 10 min pre-incubation at 40 °C to promote selective hybridization to WT regions **(Figure S3A-C)**. An amplification time of 35 min yielded strong signals for mutant templates while preventing detectable WT amplification **(Figure S3D)**. Real-time C-LAMP analysis and temperature tolerance studies (±2 °C variation) further confirmed robust WT suppression without leakage amplification, supporting the thermal stability of the LNA-clamping strategy **(Figures S4 and S5)**. DNA integrity was verified using *ACTB* as a reference control during isothermal amplification **(Figure S3E)**.

Integration of C-LAMP with ^1^O_2_-driven PEC readout was subsequently optimized using b-CPs specific for *KRAS* G12V and gDNA from SW620 (G12V/G12V) cell line and NTC samples **(Figure 2A)**. Key parameters, including C-LAMP product input volume, b-CP concentration, DP concentration and hybridization time, were systematically evaluated **(Figure 2B-E)**. The highest SW620/NTC photocurrent ratio was achieved using 5 µL of C-LAMP product, 100 nM b-CP, 50 nM DP and 15 min hybridization. Additional variables such as magnetic bead volume, amplicon denaturation and hybridization buffer composition were also examined, where 5 µL of Strep-MBs and denaturation markedly improved signal discrimination **(Figure S6A-B)**. While increased ionic strength enhanced overall PEC signal intensity, it also elevated background currents in NTC samples, consistent with reduced mismatch discrimination under high-salt concentrations (You et al. 2006). Therefore, the original hybridization buffer was retained to maintain optimal SW620-to-NTC performance **(Figure S6C)**. Collectively, these optimization studies established a reproducible workflow combining WT suppression, stable hybridization of mutant C-LAMP products and sensitive photocurrent generation using gDNA.

**Figure 2.**
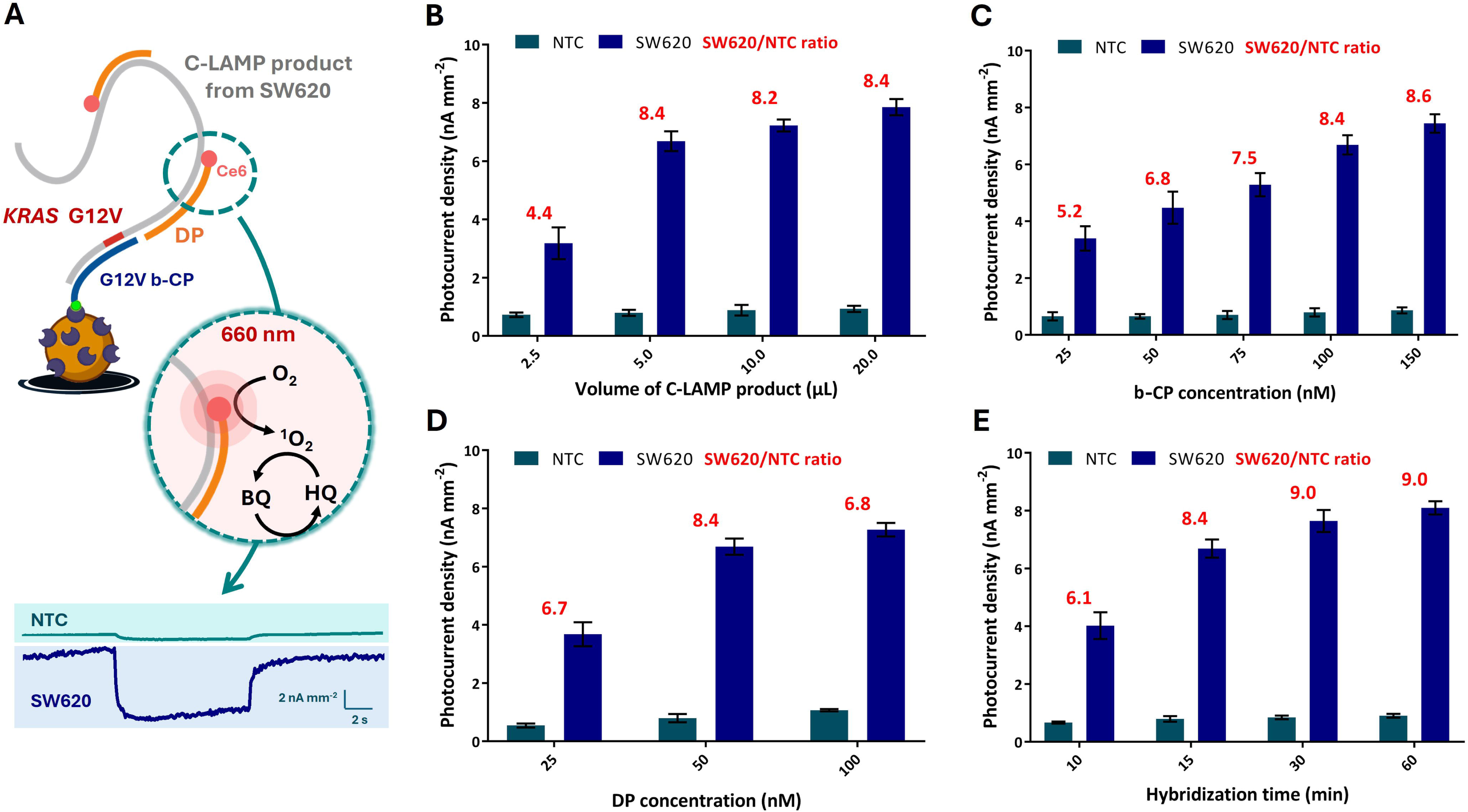
Optimization of the C-LAMP/^1^O_2_-driven PEC platform using gDNA from SW620 (G12V/G12V) and NTC, with b-CPs specific for *KRAS* G12V detection. **(A)** Schematic representation of C-LAMP product hybridization and ^1^O_2_-driven PEC readout. Optimization of key parameters: **(B)** input volume of C-LAMP products, **(C)** b-CP concentration, **(D)** DP concentration, and **(E)** hybridization time. Photocurrent ratios between SW620 and NTC samples are indicated in red. Error bars represent the standard deviation of the mean (n = 3).

### 2.3 Analytical performance in cancer cell lines

The analytical performance of the C-LAMP/^1^O_2_-driven PEC platform was evaluated using gDNA extracted from HT-29 and SW620 cancer cell lines **(Table S1)**. Conventional LAMP and C-LAMP reactions were performed under identical conditions to assess the impact of WT suppression **(Figure 3A-C)**. In standard LAMP, both WT and mutant templates were amplified and signal discrimination relied primarily on the hybridization selectivity of the b-CP, yielding modest photocurrent ratios (∼1.8). In contrast, incorporation of LNA clamp probes in C-LAMP efficiently suppressed WT amplification while maintaining strong mutant signals, increasing the discrimination ratio to ∼6.2. This threefold improvement highlights the critical role of WT blocking in improving SNV resolution in gDNA from cell lines.

**Figure 3.**
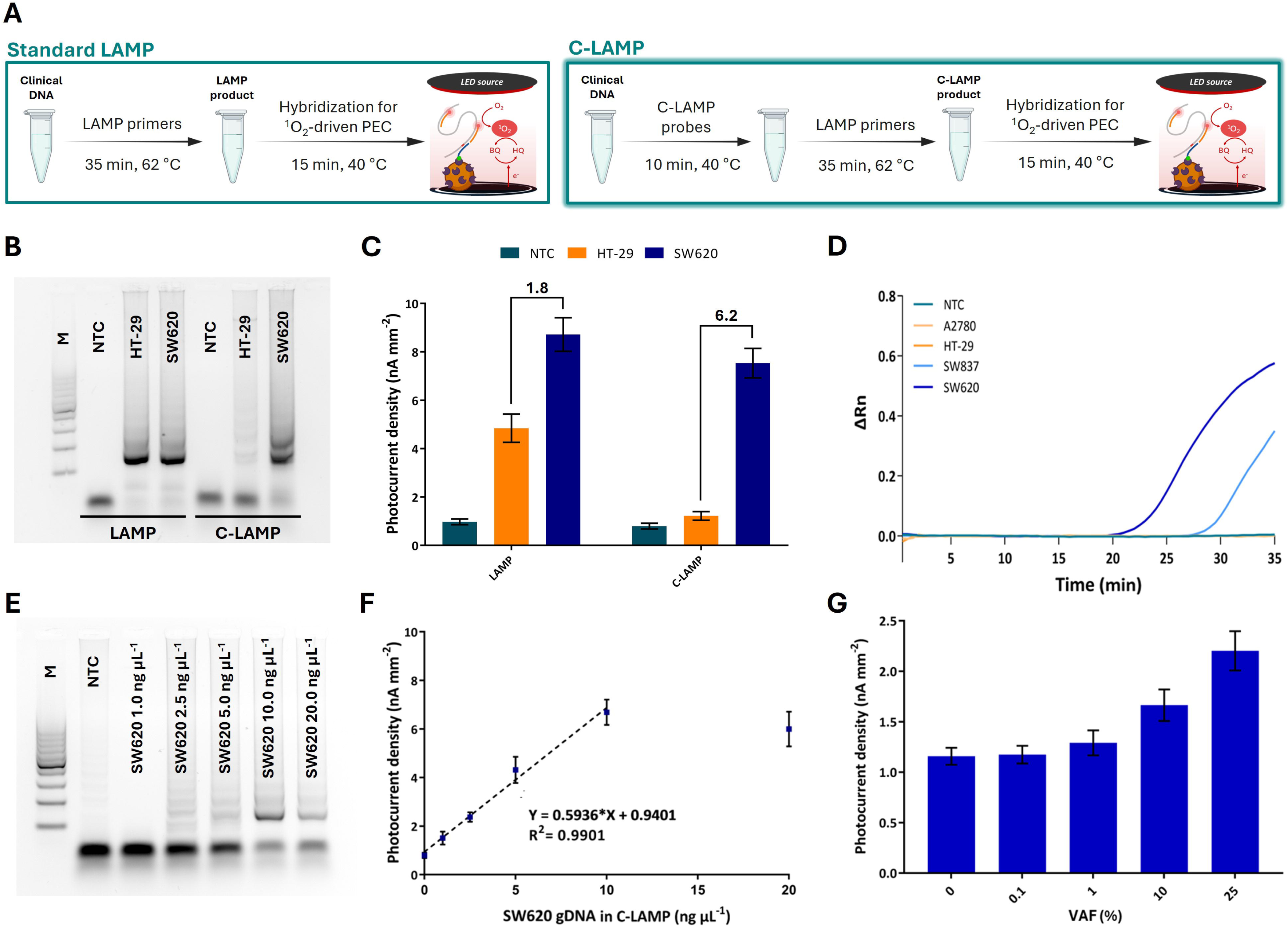
Analytical performance of the C-LAMP/^1^O_2_-driven PEC platform in gDNA from cell lines. **(A)** Schematic comparison of conventional LAMP and C-LAMP workflows. **(B)** Agarose gel electrophoresis of C-LAMP products from NTC, WT (HT-29) and mutant (SW620). **(C)** ^1^O_2_-driven PEC responses for LAMP and C-LAMP, demonstrating enhanced mutant/WT discrimination with clamp-mediated WT suppression. Photocurrent density ratios (SW620/HT-29) are indicated above the bars. **(D)** Real-time C-LAMP amplification curves obtained under optimized conditions. **(E)** Agarose gel electrophoresis of C-LAMP products from serial dilutions of SW620 gDNA. **(F)** Corresponding ^1^O_2_-driven PEC calibration plot showing linear response between 1-10 ng µL^-1^ and exhibiting a LOD of 35 copies µL^-1^ (58 aM) based on ddPCR quantification. NTC samples from the C-LAMP reaction was used as blanks. **(G)** Photocurrent densities obtained from mixtures of SW620 and HT-29 gDNA at increasing VAF while maintaining constant total gDNA concentration (10 ng µL^-1^), estimating a minimum detectable VAF of 4.8%. M: 100-1,000 bp DNA ladder. Error bars represent the standard deviation of the mean (n = 3).

To further validate amplification selectivity under optimized conditions, real-time C-LAMP experiments were performed **(Figure 3D)**. Robust amplification kinetics were observed exclusively for mutant templates, whereas WT and NTC samples remained at baseline throughout the reaction window used for isothermal amplification. These results confirm the absence of detectable non-specific products and support the specificity of the LNA-clamping strategy under the optimized assay conditions.

Analytical sensitivity was assessed using serial dilutions of SW620 gDNA. Agarose gel electrophoresis confirmed amplification down to 2.5 ng µL^-1^, whereas ^1^O_2_-driven PEC detection enabled quantitative readout down to 1 ng µL^-1^ **(Figure 3E-F)**. The calibration plot exhibited linearity between 1 and 10 ng µL^-1^ (R^2^ = 0.9901). The limit of detection (LOD), calculated according to Equation 2 (Supplementary Information) using gDNA target-free control as blank, was 0.54 ng µL^-1^. The molar concentration was estimated based on droplet digital PCR (ddPCR) quantification, providing 5,666 *KRAS* copies per 100 ng of the analyzed SW620 gDNA **(Figure S7)**. At the LOD of the platform, this corresponds to 35 copies µL^-1^ or 58 aM using Equation 4 (Supplementary Information).

To determine the minimum detectable VAF, spiking experiments were performed by mixing SW620 (mutant) and HT-29 (WT) gDNA at defined ratios while maintaining a constant total DNA input of 10 ng µL^-1^, showing increased photocurrent density at higher VAFs **(Figure 3G)**. Using linear regression within the responsive range **(Figure S8A)** and applying the LOD criterion (Equation 2), the minimum detectable VAF was estimated to be 4.8% under optimized conditions. Effective WT suppression and quantitative response were maintained across heterogeneous mutant/WT mixtures, supporting the potential applicability of the biosensing platform to clinical samples with 5-25% VAF.

Finally, a dedicated reproducibility study at 10 ng µL^-1^ of gDNA using Equation 3 (Supplementary Information) and independently prepared electrodes and bead suspensions **(Figure S8B)** yielded intra-/inter-assay RSDs of 9.8%/11.9% for HT-29 and 6.0%/8.1% for SW620 (n = 8), confirming robust fabrication and measurement reproducibility of the C-LAMP/^1^O_2_-driven PEC platform.

### 2.4 Specificity and variant discrimination

No false-positive signals were observed in WT samples (HT-29, A2780), confirming effective clamp-mediated suppression of WT amplification and absence of detectable non-specific PEC responses **(Figure 4A)**. In contrast, mutant cell lines SW837 (G12C/WT) and SW620 (G12V/G12V) produced strong amplification bands and corresponding photocurrent signals, consistent with their genotypic status. These findings were consistent with real-time C-LAMP experiments **(Figure 3D)**, which confirmed robust wild-type suppression under optimized conditions. In addition, receiver operating characteristic (ROC) analysis was performed to evaluate classification performance **(Figure 4B)**. A threshold of 3.2 nA mm^−2^ was estimated using Youden’s J statistic (see Suplementary Information) and enabled complete discrimination between WT and mutant samples, yielding an area under the curve (AUC) of 1 within the evaluated dataset.

**Figure 4.**
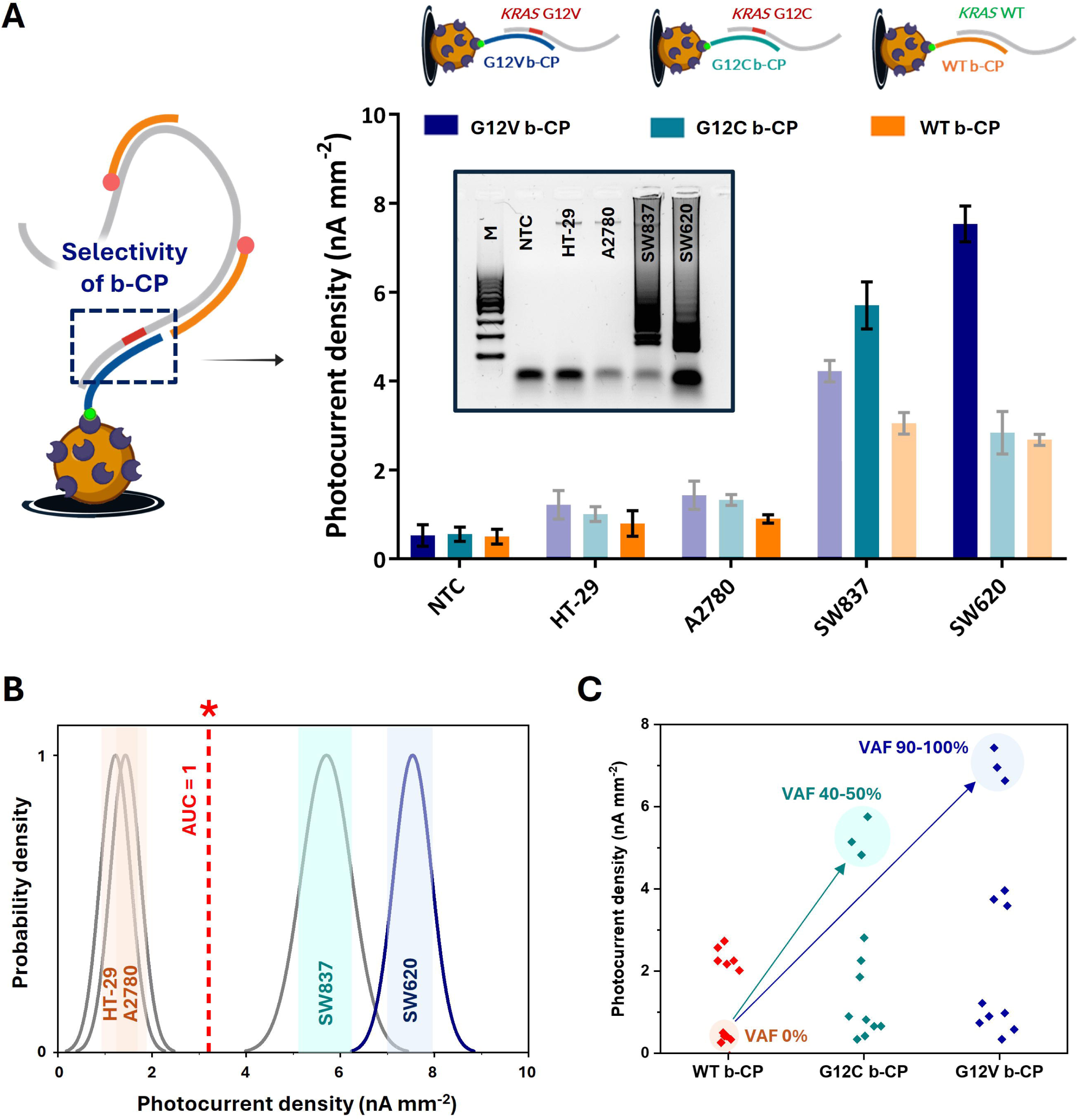
Specificity of the C-LAMP/^1^O_2_-driven PEC platform. **(A)** Photocurrent densities obtained from NTC, WT (HT-29, A2780) and mutant (SW837, G12C/WT; SW620, G12V/G12V) cell lines mutation-specific b-CPs targeting *KRAS* G12V (dark blue), G12C (turquoise) and WT sequences (orange). The inset shows agarose gel electrophoresis of C-LAMP products confirming selective amplification of mutant templates under optimized conditions. M: 100-1,000 bp DNA ladder. **(B)** ROC-derived probability distributions with a cut-off value of 3.2 nA mm^-2^, enabling complete separation between WT and mutants samples (red dashed line, *) within the evaluated dataset, yielding an AUC of 1. **(C)** Correlation between photocurrent densities and b-CPs for each cell line, demonstrating that PEC output reflects genotype-dependent intensity and that perfectly matched probe-target pairs generate the highest PEC signal response. Error bars represent the standard deviation of the mean (n = 3).

Genotyping capability was further examined using mutation-specific b-CPs targeting *KRAS* WT, G12C and G12V sequences. Perfectly matched probe–target pairs generated the highest photocurrent densities, whereas single-mismatch combinations resulted in reduced but still detectable PEC signals **(Figure 4A)**. In particular, partial cross-reactivity was observed between the G12V b-CP and G12C amplicons, consistent with the presence of a single nucleotide difference at codon 12. Importantly, for each mutant cell line (SW837 and SW620), the photocurrent obtained with the perfectly matched capture probe was substantially higher than that generated by non-cognate capture probes **(Figure 4C)**, preserving discriminatory capacity. WT-only samples remained at baseline under all tested conditions, indicating that the observed cross-reactivity reflects hybridization effects rather than amplification leakage or false-positive generation.

From a clinical perspective, activating *KRAS* mutations such as G12V and G12C are typically mutually exclusive within a single tumor. Therefore, reliable identification of any activating *KRAS* mutation is often sufficient for therapeutic stratification, particularly in the context of anti-EGFR treatment decisions. While precise differentiation between closely related variants may require further probe refinement to minimize residual cross-reactivity, the present platform provides robust discrimination between WT and mutant *KRAS* sequences and enables differentiation of closely related variants under optimized conditions.

### 2.5 Clinical validation in patient-derived tissues

The translational applicability of the C-LAMP/^1^O_2_-driven PEC platform was evaluated using gDNA extracted from fresh frozen tissue (FFT) samples obtained from NSCLC patients **(Table S2)**. Under optimized conditions, C-LAMP selectively amplified mutant alleles while suppressing WT templates, as confirmed by agarose gel electrophoresis **(Figure 5A)**. WT samples (e.g., FFT-2 and FFT-8) showed no visible amplification bands, whereas mutant samples as FFT-11 (*KRAS* G12C) and FFT-15 (*KRAS* G12V) produced clear amplification products.

**Figure 5.**
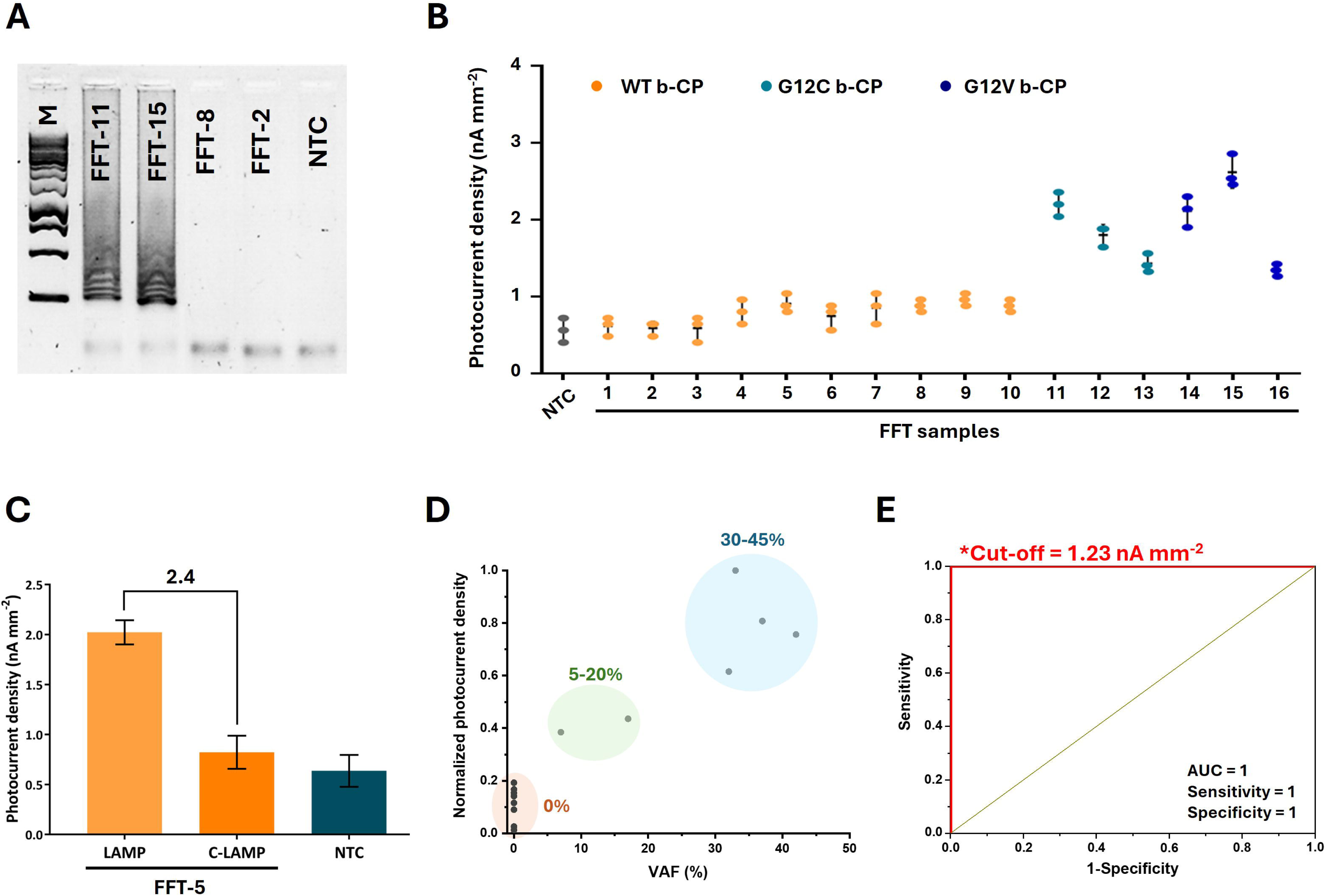
Clinical validation of the C-LAMP/^1^O_2_-driven PEC platform in FFT samples. **(A)** Agarose gel electrophoresis of C-LAMP products obtained from representative WT controls (FFT-2 and FFT-8) and *KRAS*-mutant patients carrying G12C (FFT-11) and G12V (FFT-15). M: 100-1,000 bp DNA ladder. **(B)** Photocurrent densities obtained with b-CPs specific for WT (orange), G12C (turquoise) and G12V (dark blue), confirming genotype-specific detection depending on probe complementarity. **(C)** Comparison of conventional LAMP and C-LAMP in a representative WT sample (FFT-5), confirming efficient WT suppression upon inclusion of LNA clamp probes during amplification. Error bars represent the standard deviation of the mean (n = 3). **(D)** Correlation between normalized photocurrent densities and VAF values determined by ddPCR. **(E)** ROC analysis for discrimination between WT and mutant FFT samples with ROC-defined threshold (*Cut-off) enabling complete separation of WT and mutant samples within the evaluated cohort (n = 16).

^1^O_2_-driven PEC readouts were consistent with these results: WT samples generated background-level photocurrents, while mutant tissues yielded genotype-specific responses depending on the b-CP employed **(Figure 5B)**. Specifically, FFT-11 to FFT-13 (G12C) responded selectively to the G12C b-CP, whereas FFT-14 to FFT-16 (G12V) responded to the G12V b-CP. Comparison between conventional LAMP and C-LAMP in a WT sample (FFT-5) revealed a more than twofold reduction in photocurrent when LNA clamp probes were included, confirming effective WT blocking during isothermal amplification **(Figure 5C)**.

VAF values determined by ddPCR **(Figures S9-S12 and Table S2)** were used as quantitative references for correlation with normalized photocurrent densities. As shown in **Figure 5D**, a positive correlation was observed using the C-LAMP/^1^O_2_-driven PEC platform in the analysis of FFT samples, indicating that PEC output reflects allelic burden across clinically relevant VAF ranges (approximately 5-40% in the evaluated cohort). Notably, FFT samples with VAF values as low as 5-20% were distinguishable from WT tissues.

ROC analysis was performed to evaluate diagnostic classification performance. Within the evaluated cohort (n = 16), complete discrimination was achieved between WT and mutant FFT samples, offering a cut-off value of 1.23 nA mm^-2^ **(Figure 5E)**. While larger clinical studies are required to establish definitive diagnostic performance metrics, these preliminary results demonstrate full concordance with gold-standard ddPCR in the analyzed samples and support the translational potential of the C-LAMP/^1^O_2_-driven PEC platform for decentralized *KRAS* mutation analysis.

## 3. Discussion

Recent advances in pan-*KRAS* inhibitors underscore the growing importance of comprehensive *KRAS* genotyping in clinical oncology (Choate et al. 2024), highlighting the need for analytical platforms capable of selectively identifying activating mutations within heterogeneous DNA backgrounds. While NGS and ddPCR remain reference standards for *KRAS* mutation analysis, their implementation in decentralized or resource-limited settings is constrained by instrumentation requirements, workflow complexity and the need for specialized infrastructure.

Recent (photo)electrochemical approaches, frequently coupled with isothermal amplification, have been explored for *KRAS* mutation detection **(Table S3)**. Although these approaches demonstrate notable analytical sensitivity, many rely on enzymatic reporters, involve complex or time-consuming workflows and, critically, lack validation in patient-derived samples. Most LAMP-based *KRAS* assays employ fluorescent or colorimetric readouts combined with PNA-modified primers or ligation-based designs (Fu et al. 2019; Islam et al. 2025; Itonaga et al. 2016; Mirlohi et al. 2024). Because mutant alleles may constitute only a small fraction of total DNA, simultaneous enrichment of mutant sequences and suppression of WT amplification are essential to achieve robust discrimination. While PNA clamps offer strong binding affinity and mismatch discrimination, their synthesis cost and solubility limitations can complicate routine implementation(Ondraskova et al. 2023). Moreover, optical detection systems remain relatively costly and less compatible with decentralized testing than (photo)electrochemical readouts. In contrast, the present approach introduces a practical alternative based on commercially available, water-soluble LNA-modified oligonucleotides that act both as allele-specific blockers during amplification and as capture probes for SNV recognition in the PEC transduction stage, thereby providing a scalable and clinically relevant route toward decentralized testing.

C-LAMP effectively mitigates WT interference, which has historically limited the selectivity of isothermal amplification methods. The dual implementation of LNA chemistry constitutes a key molecular design feature underlying the platform’s specificity. In the amplification stage, LNA incorporation within the clamp probes increases duplex stability and mismatch discrimination, enabling complete WT suppression under isothermal conditions. In the PEC transduction stage, LNA substitutions in the capture probes enhance hybridization affinity toward mutant amplicons while minimizing off-target binding. Nevertheless, limited cross-reactivity between closely related variants (e.g., G12C and G12V) was observed, consistent with the high sequence homology (>93%) shared among b-CPs and the inherent thermodynamic constraints of single-mismatch discrimination under isothermal conditions. Importantly, such partial cross-reactivity did not result in false-positive classification of WT samples and therefore does not compromise mutant-versus-WT discrimination, which represents the primary clinical requirement for diagnosis and many therapeutic decision pathways.

The enzyme-free ^1^O_2_-driven PEC readout further contributes to assay robustness by avoiding enzymatic reporters such as horseradish peroxidase (HRP), which require external substrates to generate electrochemical signals and controlled storage conditions. Photosensitizers such as Ce6 generate ^1^O_2_ directly under visible-light excitation, promoting HQ/BQ redox cycling without enzymatic intermediates. This photochemical transduction simplifies reagent handling, eliminates instability associated with enzyme storage and reduces per-test cost, key considerations for decentralized implementations.

Comparison with state-of-the-art diagnostic methods **(Table S4)** highlights the complementary positioning of the C-LAMP/¹O₂-driven PEC platform. Whereas ddPCR and NGS provide absolute quantification and base-level sequence resolution, the proposed system offers a simplified hardware configuration requiring a constant-temperature heater, a low-power light source and a portable potentiostat. With total amplification and detection times close to one hour, the workflow may serve as a rapid screening or triage tool in molecular oncology settings. Importantly, the positive correlation between photocurrents and VAF values in both cell lines and patient-derived samples indicates that PEC output reflects mutation burden, enabling semi-quantitative assessment without sequencing. Moreover, although the clinical dataset is limited (n = 16), the absence of misclassification within this cohort suggests high discriminative capability for potential adoption in decentralized testings.

Regarding potential interference effects from complex biological matrices, it is important to note that the majority of experiments were conducted using purified gDNA extracted from clinically characterized samples. Such DNA extracts reflect standard diagnostic workflows and avoid reliance on synthetic constructs, thereby providing a clinically relevant evaluation environment. The successful discrimination of *KRAS* mutation status under these conditions supports the practical applicability of the proposed assay within routine genomic testing frameworks.

Several aspects warrant further refinement. Hybridization kinetics between C-LAMP products and capture probes could be optimized to shorten assay time, and expansion to additional *KRAS* variants (e.g., G12D, G13D) will require probe design and validation. Although partial cross-reactivity was observed between certain codon 12 variants, activating *KRAS* mutations are typically mutually exclusive within individual tumors, reducing the likelihood of misclassification in routine screening contexts. Nevertheless, applications requiring strict mutation-specific stratification (e.g., allele-targeted inhibitor selection) would benefit from further probe optimization to enhance discrimination stringency.

Finally, while the present study relies on pre-extracted gDNA in accordance with standard molecular diagnostic workflows (such as ddPCR), integration with simplified sample-preparation modules or microfluidic cartridges may further streamline the assay architecture. Larger clinical validation studies will be necessary to establish definitive diagnostic performance metrics. Within the evaluated cohort, however, full concordance with ddPCR was achieved, supporting the translational potential of this C-LAMP/PEC platform.

Overall, the integration of clamp-mediated isothermal amplification with enzyme-free ^1^O_2_-driven PEC detection establishes a stable and analytically robust biosensing framework. By merging molecular selectivity with light-driven electrochemical transduction, this approach provides a versatile strategy for mutation detection that may be extended to other actionable oncogenic variants and nucleic-acid biomarkers.

## 4. Experimental/methods section

### 4.1 Instrumentation and electrode configuration

^1^O_2_-driven PEC measurements were performed at room temperature using a PalmSens4 potentiostat operated with PSTrace 5.9 software (PalmSens, The Netherlands). Screen-printed carbon electrodes (DRP-110, ф = 0.4 cm) served as transducers and were connected through a DSC box connector (Metrohm DropSens, Spain). Illumination was provided by a 660 nm LED integrated into a pE-4000 system (CoolLED, UK) synchronized with the potentiostat via the digital I/O lines of PSTrace. The LED power (30 mW) was calibrated using a PM100D optical power meter (Thorlabs, USA). Real-time C-LAMP experiments were conducted on the QuantStudio 5 device with direct analysis by the QuantStudio software (Thermo Fisher Scientific, USA).

### 4.2 Oligonucleotide design and sequences

The complete list of oligonucleotide sequences and terminal modifications is provided in **Table S4**. LAMP primers (F3, B3, FIP and BIP) were designed using PrimerExplorer V.5 (Eiken Chemical Co., Japan) and synthesized by Integrated DNA Technologies (USA). LNA-modified clamp probes (CL1 and CL2) and capture probes (a-CPs and b-CPs), were obtained from the same supplier. DPs were synthesized and purified by reverse-phase HPLC, and subsequently labelled with Ce6 as the photosensitizer (Eurogentec, Belgium). All oligonucleotides were dissolved in nuclease-free PCR-grade water (VWR, Avantor^®^, USA) and stored at −20 °C until use.

The initial C-LAMP primer set was designed to amplify multiple *KRAS* SNVs without targeting a specific mutation, with downstream discrimination achieved through hybridization with mutation-specific capture probes. However, this design exhibited reduced amplification efficiency for WT templates, primarily due to the backward inner primer (BIP.V2; **Table S4**). To address this limitation, alternative BIP sequences (BIP and BIP.V3) were designed to anneal closer to the mutation hotspot. The original primer contained only a single guanine overlap at the *KRAS* mutation site, whereas the new sequences spanned both codons, either with (GGTGGCG) or without (GGTGGC) additional nucleotides. The most consistent performance was obtained with the latter design, in which the BIP primer hybridized across both codons without consecutive mismatches. This optimized sequence (BIP) was selected for the final C-LAMP protocol.

### 4.3 Cancer cell lines

Cancer cell lines used as models for *KRAS* mutation detection are listed in **Table S1**. The selected cell lines were chosen to represent homozygous (WT and mutant) and heterozygous *KRAS* mutation backgrounds, enabling systematic evaluation of WT suppression and mutation-specific amplification across different allelic compositions. HT-29 and A2780 (WT/WT), SW837 (G12C/WT) and SW620 (G12V/G12V) were cultured in Dulbecco’s Modified Eagle’s Medium (DMEM) supplemented with 10% fetal bovine serum, 1% penicillin-streptomycin and 1% sodium pyruvate under standard conditions (37 °C, 5% CO_2_ and 100% humidity). Cells were collected by scraping, centrifuged (1,500 rpm, 5 min) and the resulting pellets stored at − 80 °C until use. gDNA was extracted using the Tissue DNA Preparation Column Kit (Jena Bioscience, Germany) according to the manufacturer’s protocol. DNA purity was assessed by NanoDrop UV-Vis spectrophotometry (Thermo Fisher Scientific, USA) and concentrations were determined using a Qubit 4 fluorometer (Invitrogen, USA) with the dsDNA HS Assay Kit.

*KRAS* amplicons were sequenced using the MinION platform (Oxford Nanopore Technologies, UK) with primers listed in **Table S4** (ddPCR Fwd and SEQ Rev). Amplicons were generated with LongAmp^®^ Hot Start Taq 2X Master Mix (New England Biolabs, USA), purified with AMPure XP magnetic beads (Beckman Coulter, USA) and barcoded using the Rapid Barcoding Kit 14 (SQK-RBK114.24, Oxford Nanopore) following the manufacturer’s instructions. Libraries were cleaned with AMPure XP beads, quantified, pooled and loaded onto a MinION flow cell (R10.4.1). Sequencing was performed using MinKNOW software; basecalling and demultiplexing were conducted in Guppy (high-accuracy mode). Reads were aligned to the *KRAS* reference sequence ((NCBI) 2025) using EPI2ME and IGV platforms. Nucleotide substitutions detected at codons 12 and 13 matched the expected genotypes for each cell line **(Table S1)**.

Droplet digital PCR (ddPCR) was performed on SW620 gDNA using the QX200 AutoDG system (Bio-Rad, USA) with *KRAS*-specific primers **(Table S6)**. Each 22 µL reaction contained 1×QX200™ EvaGreen Supermix; 0.45 µM ddPCR Fwd and ddPCR Rev primers, PCR-grade water, and 25, 50 or 100 ng of gDNA. Droplets were generated using the Automated Droplet Generator and thermal cycling was conducted as follows: 94 °C for 10 min; 35 cycles of 94 °C for 30 s; 62 °C for 30 s; 72 °C for 40 s; final extension at 72 °C for 10 min; hold at 4 °C. Fluorescence was measured with the QX200 Droplet Reader and data were processed using QuantaSoft 1.7.4.0917 software (Bio-Rad).

To determine the *KRAS* copy number in 100 ng of gDNA, copy numbers per microliter of reaction mixture were multiplied by 22 (total reaction volume, 22 µL). The 25 ng and 50 ng samples used to reduce bias caused by distribution of DNA into the droplets were normalized to 100 ng of gDNA. The mean value across replicates corresponded to 5,666 *KRAS* copies per 100 ng of SW620 gDNA, which was used to construct the SW620 calibration curve and estimate the LOD of the biosensing platform **(Figure 3F)**.

### 4.4 Patient-derived tissues

All procedures involving human material were approved by the institutional ethics committee of the University Hospital Antwerp (UZA/University of Antwerp, BUN B3002023000515) and informed consent was obtained from all participants prior to sample collection. Biobank Antwerp (ID: BE 71030031000) was also involved in this study. FFT samples, including tumor type, percentage of neoplastic cells and VAF are summarized in **Table S2**. Hematoxylin–eosin (H&E)-stained sections were prepared from each specimen and examined by a pathologist to confirm tumor content and estimate neoplastic-cell percentage.

gDNA was extracted from the remaining FFT material using the QIAamp DNA Mini Kit with a QIAshredder column (Qiagen, Germany). Briefly, pre-chopped tissue was lysed in ATL/AL buffer with Proteinase K, homogenized, incubated at 70 °C and clarified using QIAshredder columns. The lysates were combined with ethanol, transferred onto QIAamp Mini spin columns, washed with AW1 and AW2 buffers, and eluted in nuclease-free water. DNA concentration was quantified using a Qubit fluorometer (Thermo Fisher Scientific, USA).

*KRAS* mutation analysis was performed on a QX200 ddPCR system (Bio-Rad, USA). Each 20 µL reaction contained 10 µL 2× ddPCR™ Supermix for Probes (no dUTP, Bio-Rad), 1 µL of 20× *KRAS* primer/probe mix (FAM + HEX, Bio-Rad), nuclease-free water and 50 fg-130 ng of gDNA. Droplets were generated using the QX200 Automated Droplet Generator and amplified in a Veriti™ thermal cycler (Applied Biosystems, USA) under the following conditions: 95 °C for 10 min; 40 cycles of 94 °C for 30 s and 55 °C for 1 min; 98 °C for 10 min; hold at 4 °C. End-point fluorescence was recorded using the QX200 Droplet Reader and data were analyzed in multiplex mode with QuantaSoft™ software. While ddPCR allowed discrimination between WT and mutant alleles, the specific *KRAS* subtype (e.g., G12C, G12V) was determined from parallel NGS of matched FFPE tissue performed during routine clinical diagnostics. This combined workflow ensured accurate confirmation of mutation status across the FFT cohort.

### 4.5 C-LAMP protocol

C-LAMP was performed in two sequential steps: selective suppression of WT alleles followed by amplification of mutant alleles carrying SNVs. The optimized 10 µL reaction mixture contained 100 ng of gDNA (10 ng µL^-1^); 0.6 µM each of clamp probes (CL1 and CL2); 0.2 µM each of outer primers (F3 and B3); 1.6 µM each of inner primers (FIP and BIP); 1x LAMP Master Mix (STM), and nuclease-free water.

In the first step, DNA templates were incubated with clamp probes at 40 °C for 10 min to promote hybridization to WT regions, followed by rapid cooling on ice. In the second step, LAMP primers and STM were added to the clamped DNA mixture, and amplification was performed at 62 °C for 35 min. Amplicons were verified by 1.5% agarose gel electrophoresis (TBE buffer, GelRed^®^ staining) and subsequently used directly for ^1^O_2_-driven PEC measurements or stored at −20 °C until use.

### 4.6 Real-time C-LAMP protocol

The Real time C-LAMP experiments were performed using a QuantStudio™ 5 Real-Time PCR System (Applied Biosystems). The reaction mixture contained the same components as the designed C-LAMP protocol, with the exception of the amplification master mix. Instead of the conventional STM Master Mix (containing SYBR Green), an STM High-ROX Master Mix (Jena Bioscience, Germany) was used. This formulation has identical amplification components but includes ROX dye as a passive reference for fluorescence normalization.

Clamp pre-incubation was carried out for 10 min at three different temperatures (38, 40, and 42°C) to evaluate thermal tolerance. Following pre-incubation, the DNA–clamp mixture was transferred to qPCR tube strips and combined with the remaining reaction components. Real-time amplification was performed at 62 °C using 30 s acquisition intervals, with fluorescence recorded at the end of each segment for a total of 70 cycles (35 min). Cycle threshold (Ct) values were automatically calculated using QuantStudio™ Design and Analysis Software with default baseline and threshold settings.

### 4.7 Preparation of the ^1^O_2_-driven PEC platform

A 5 µL aliquot of Strep-MBs was washed twice with 200 µL of hybridization buffer on a magnetic rack (2 min per wash). Beads were then incubated with 100 µL of 100 nM b-CP solution in hybridization buffer for 10 min at 25 °C under constant agitation (950 rpm). After two additional washes, functionalized Strep-MBs were hybridized with denatured C-LAMP products. The amplicons were denatured at 95 °C for 10 min and rapidly cooled on ice before use. For hybridization, 95 µL of 50 nM DP solution was mixed with 5 µL of denatured C-LAMP products and incubated with the Strep-MB/b-CP complexes at 40 °C for 15 min with shaking (950 rpm). The resulting biofunctionalized beads were washed twice with 200 µL of measuring solution and resuspended for ^1^O_2_-driven PEC analysis.

For electrode assembly, 100 µL of 1 mM HQ in measuring solution was deposited onto a screen-printed carbon electrode, covering the three-electrode system. Biofunctionalized beads were resuspended in 10 µL of the HQ droplet, transferred onto the electrode surface and magnetically captured at the working electrode using a neodymium magnet positioned beneath the screen-printed electrode. PEC measurements followed the ^1^O_2_-mediated detection strategy(Daems et al. 2024; Shanmugam et al. 2024; Stratulat et al. 2025; Trashin et al. 2017). A potential of −0.20 V was applied to the Ag pseudo-reference electrode to reduce BQ to HQ under a light cycle of 60 s dark, 10 s illumination and 30 s dark; completing total measurement times below two minutes.

Further details on instrumentation, magnetic microbeads, reagents, and statistical analyses are provided in the Supplementary Information.

## Acknowledgements

This work was supported by the Interuniversity Special Research Fund (iBOF/23/030). Financial support from the Czech Health Czech Science Foundation (No. 25-15990S), project SALVAGE (OP JAC; reg. no. CZ.02.01.01/00/22_008/0004644) co-funded by the European Union and the State Budget of the Czech Republic, Czech Health Research Council (No. NU23J-08-00006), BBMRI.cz (No. LM2023033) and MH CZ-DRO (MMCI, 00209805) is also acknowledged. A.V. acknowledges funding from the Research Foundation-Flanders (FWO; Postdoctoral fellowship 7028-1251525N) and the Research Fund of the University of Antwerp (BOF/KP, 542800003-53370). K.D.W. acknowledges funding from the University of Antwerp (BOF/IOF/SEP) and FWO. T.V. acknowledges funding from Research Foundation-Flanders (FWO; Senior Clinical Investigator fellowship 1803723N). The authors also thank the Biobank Antwerp (ID: BE 71030031000) for providing clinical samples and the SOCan consortium for fostering collaboration and scientific discussions.

## Declarations

### Experimental ethics

All experiments were performed in compliance with institutional guidelines and in accordance with the ethical standards of the Declaration of Helsinki.

### Declaration of generative AI and AI-assisted technologies

During the preparation of this work, the authors used ChatGPT and PaperPal to improve the readability of the text. All suggestions generated by these AI-tools were carefully reviewed and edited by the authors, who take full responsibility for the final content of the article.

## Author contributions

**J.S. and A.V.:** Conceptualization, Methodology, Validation, Formal analysis, Investigation, Data curation, Visualization, Writing – Original Draft, Writing – Review & Editing. **L.M.:** Conceptualization, Methodology, Validation, Investigation, Writing – Review & Editing. **J.A.:** Validation, Investigation, Data curation, Writing – Review & Editing. **F.Z-K.:** Validation, Investigation. **S.K.:** Resources. **K.Z.:** Resources, Writing – Review & Editing. **T.V.:** Resources, Writing – Review & Editing, Supervision. **M.B. and K.D.W.:** Resources, Writing – Review & Editing, Supervision, Project administration, Funding acquisition.

## Data availability statement

All data supporting the findings of this work are available within the paper and its Supplementary Information. Additional raw data are available from the corresponding authors upon reasonable request.

---

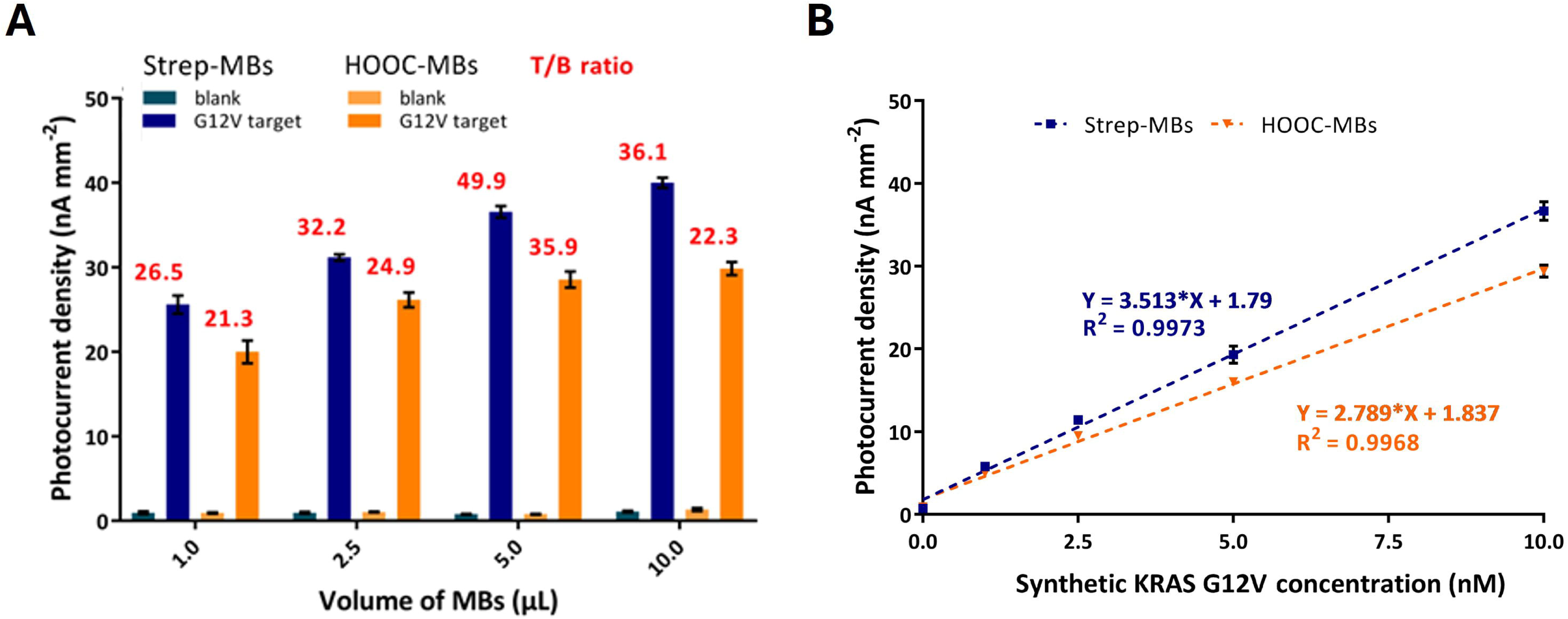

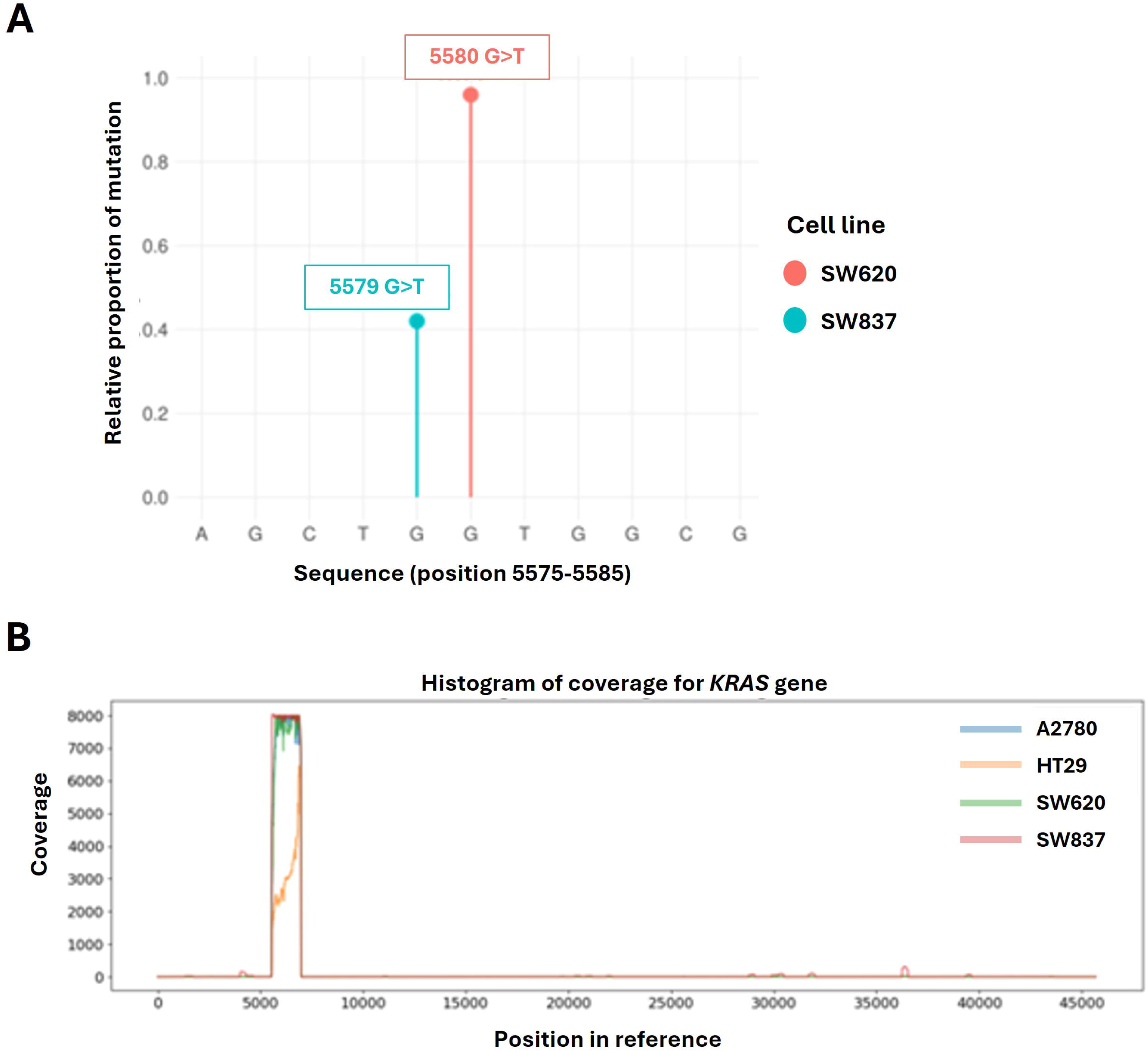

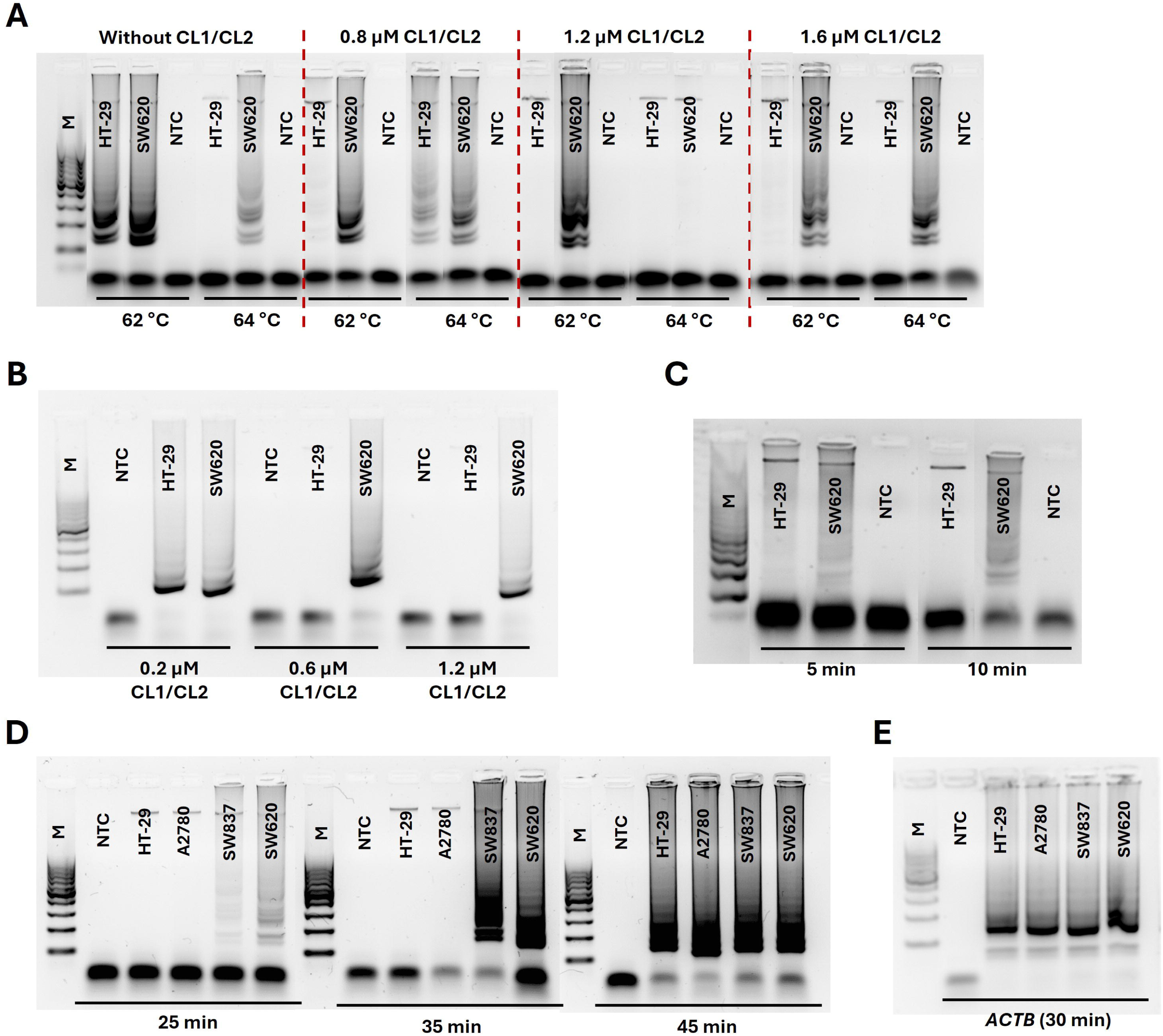

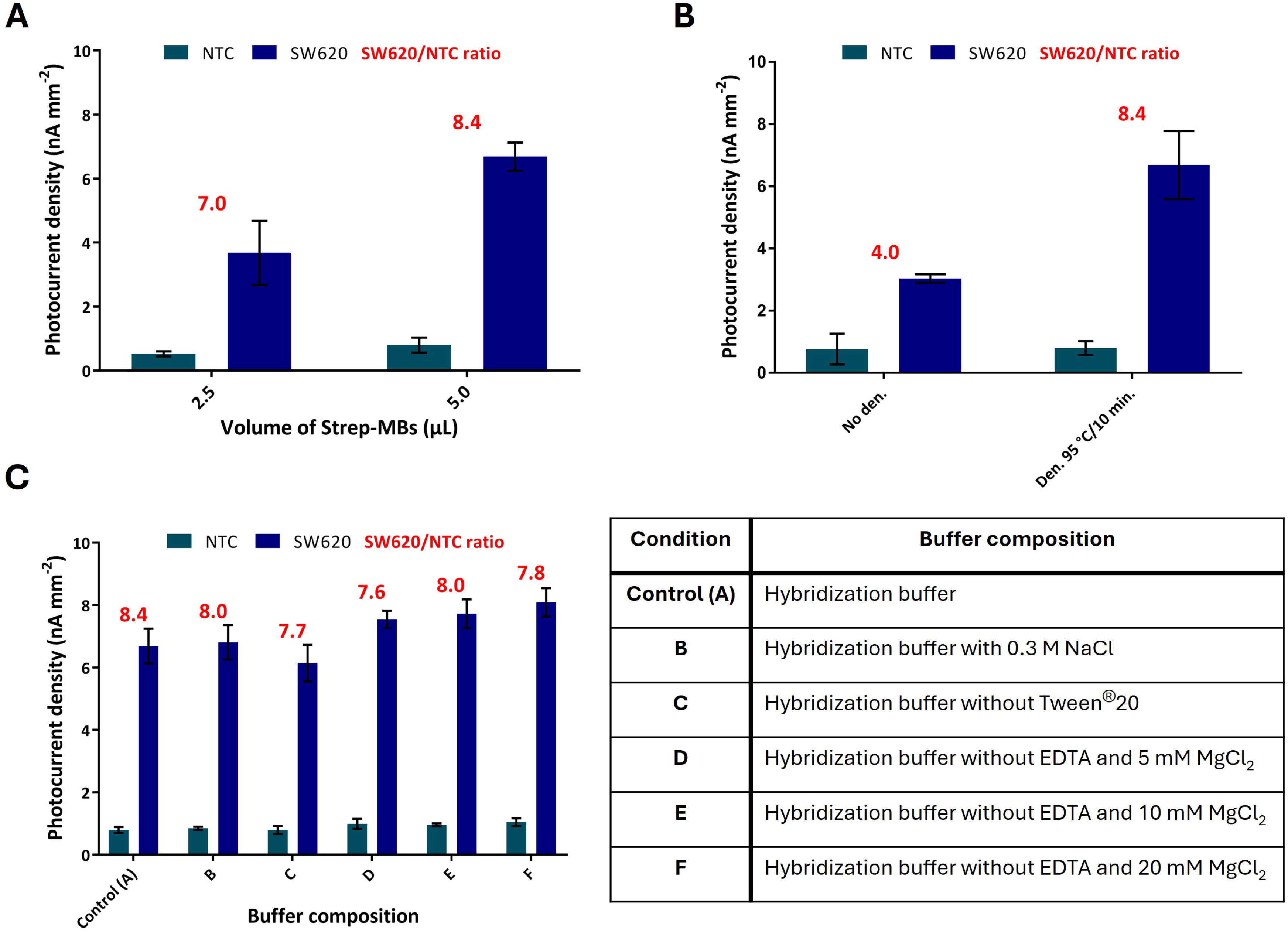

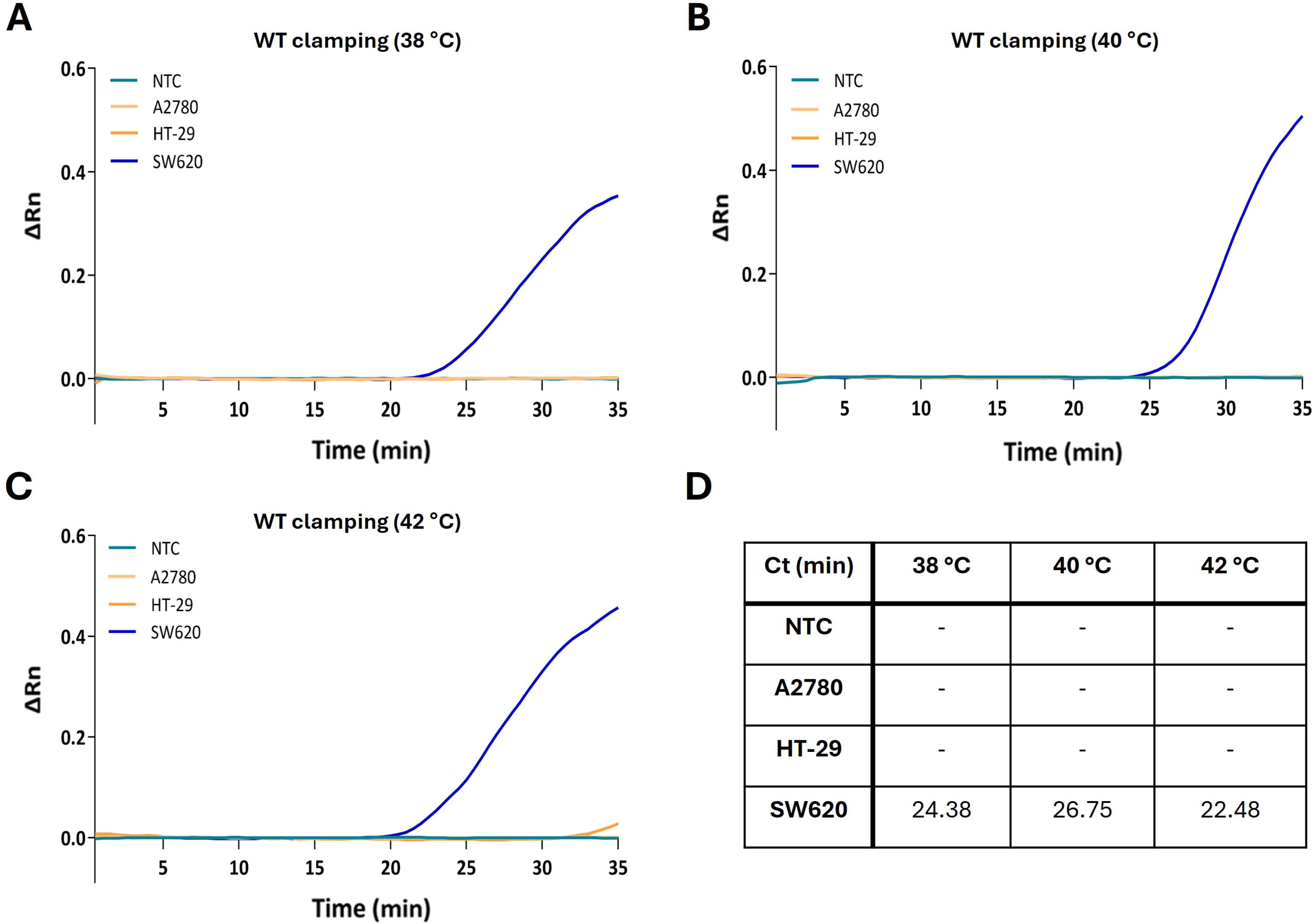

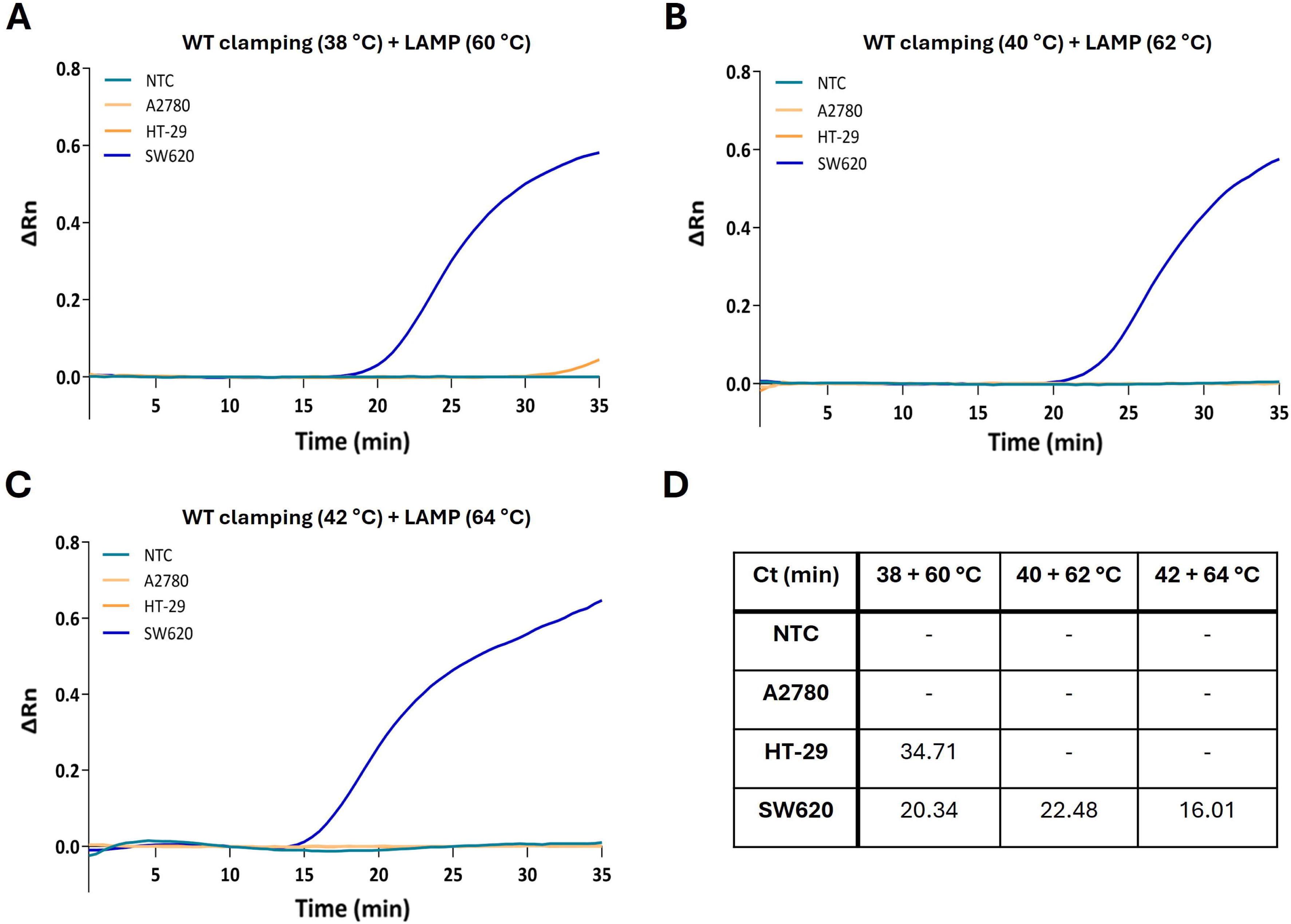

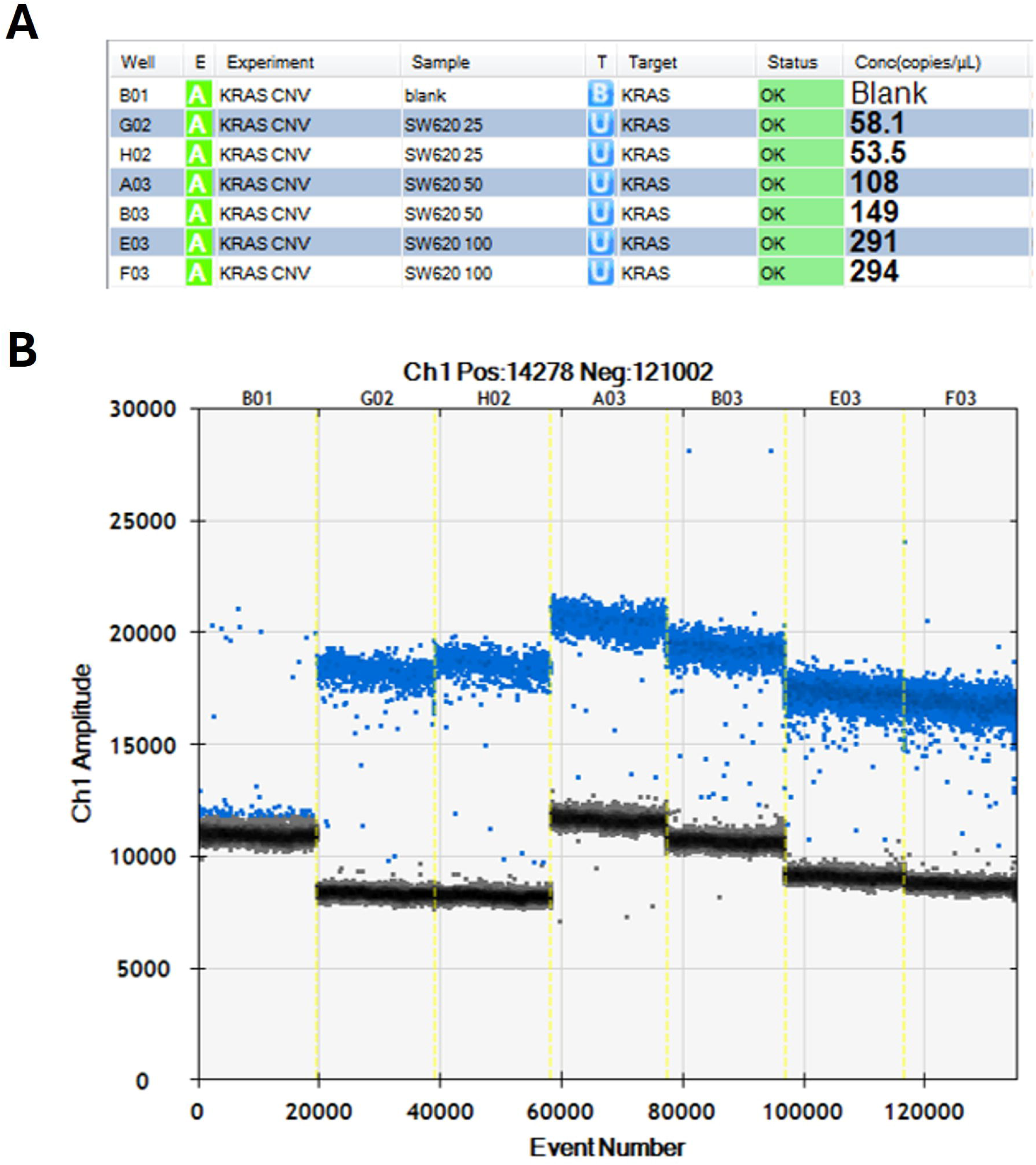

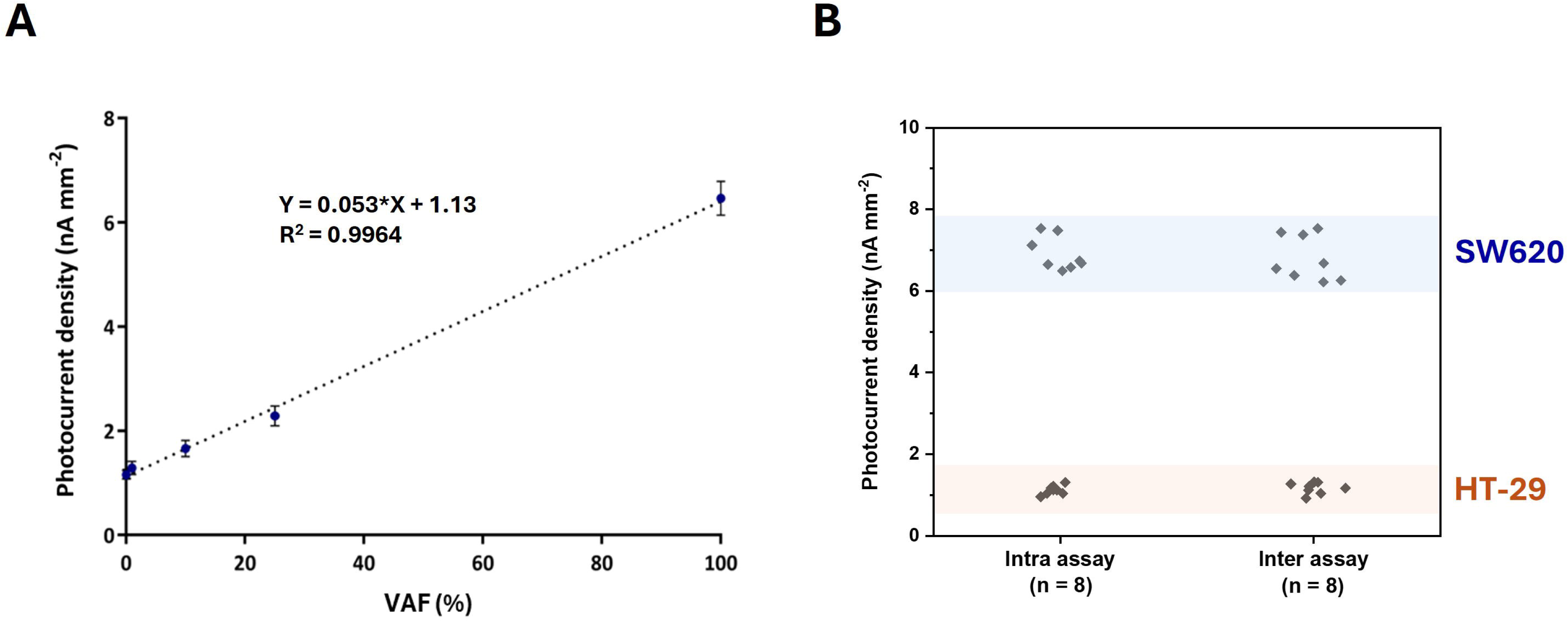

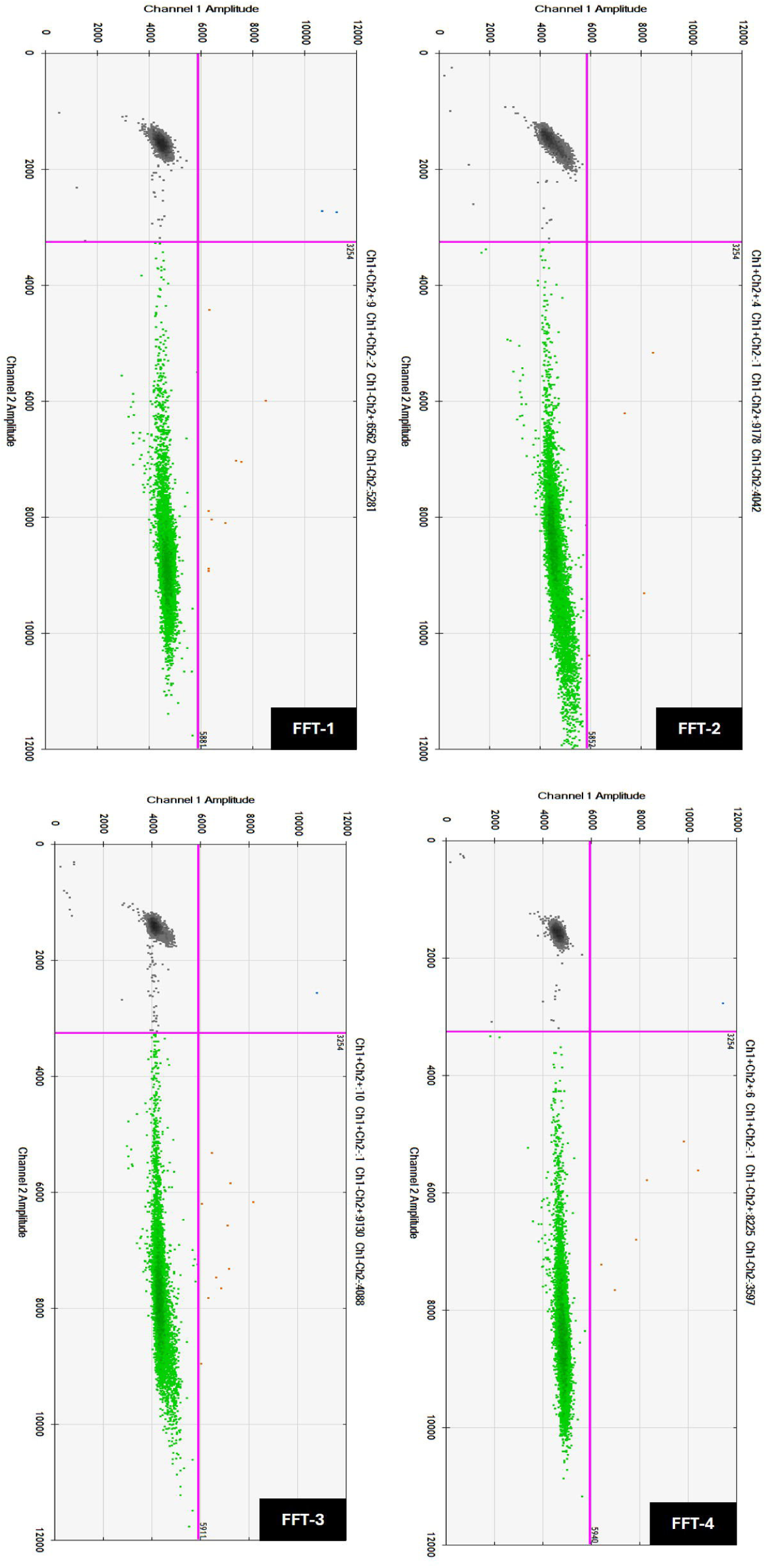

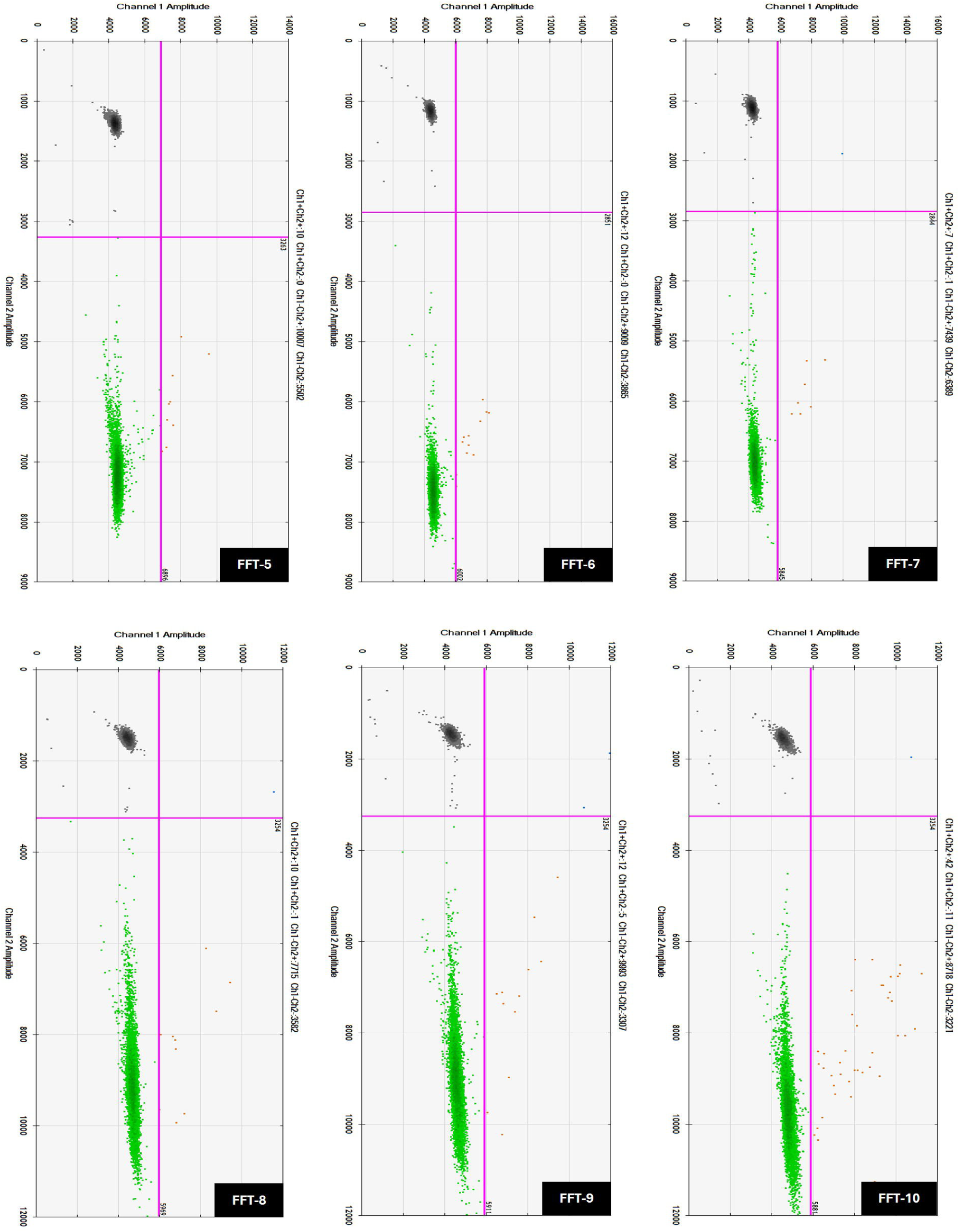

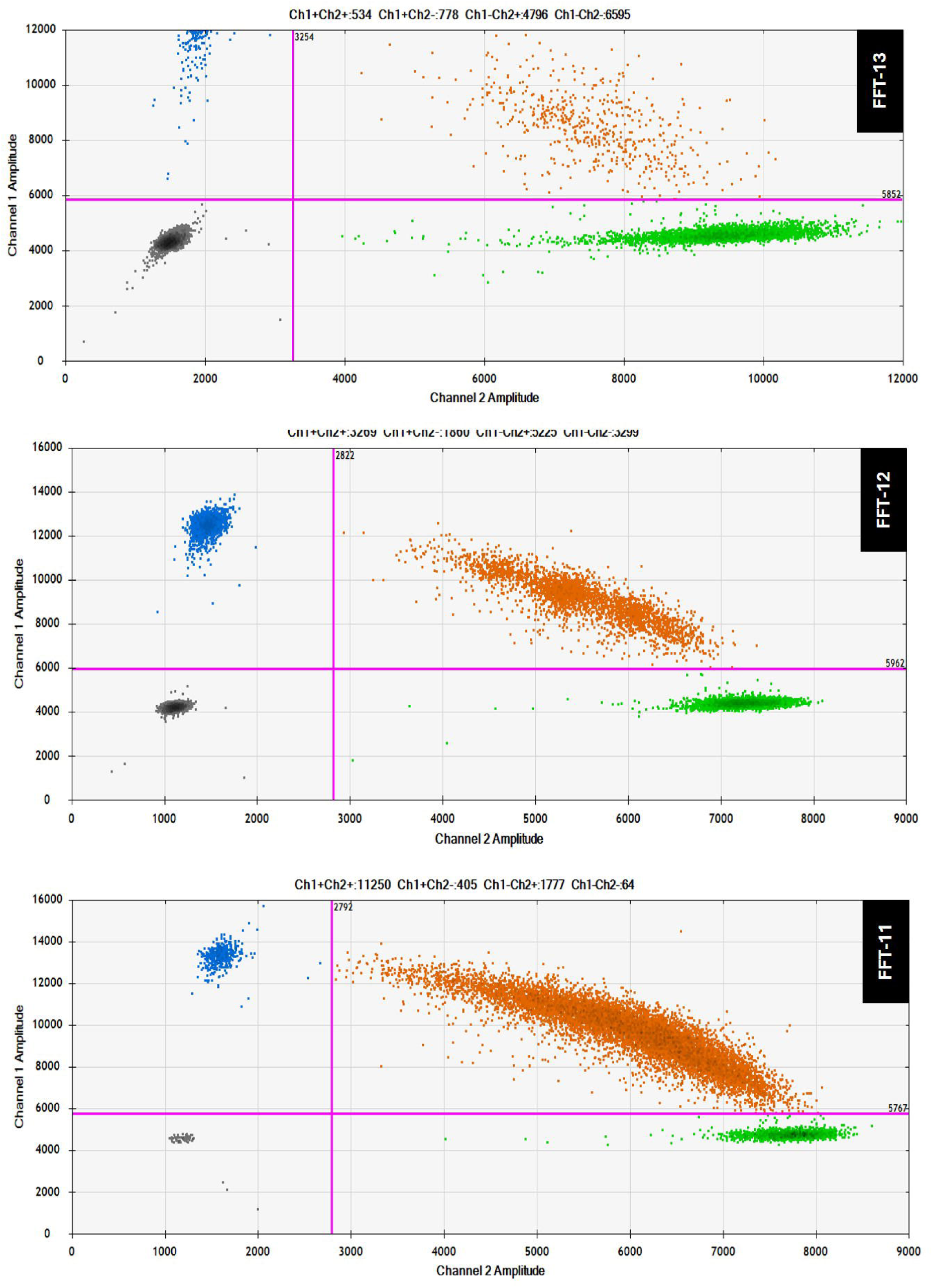

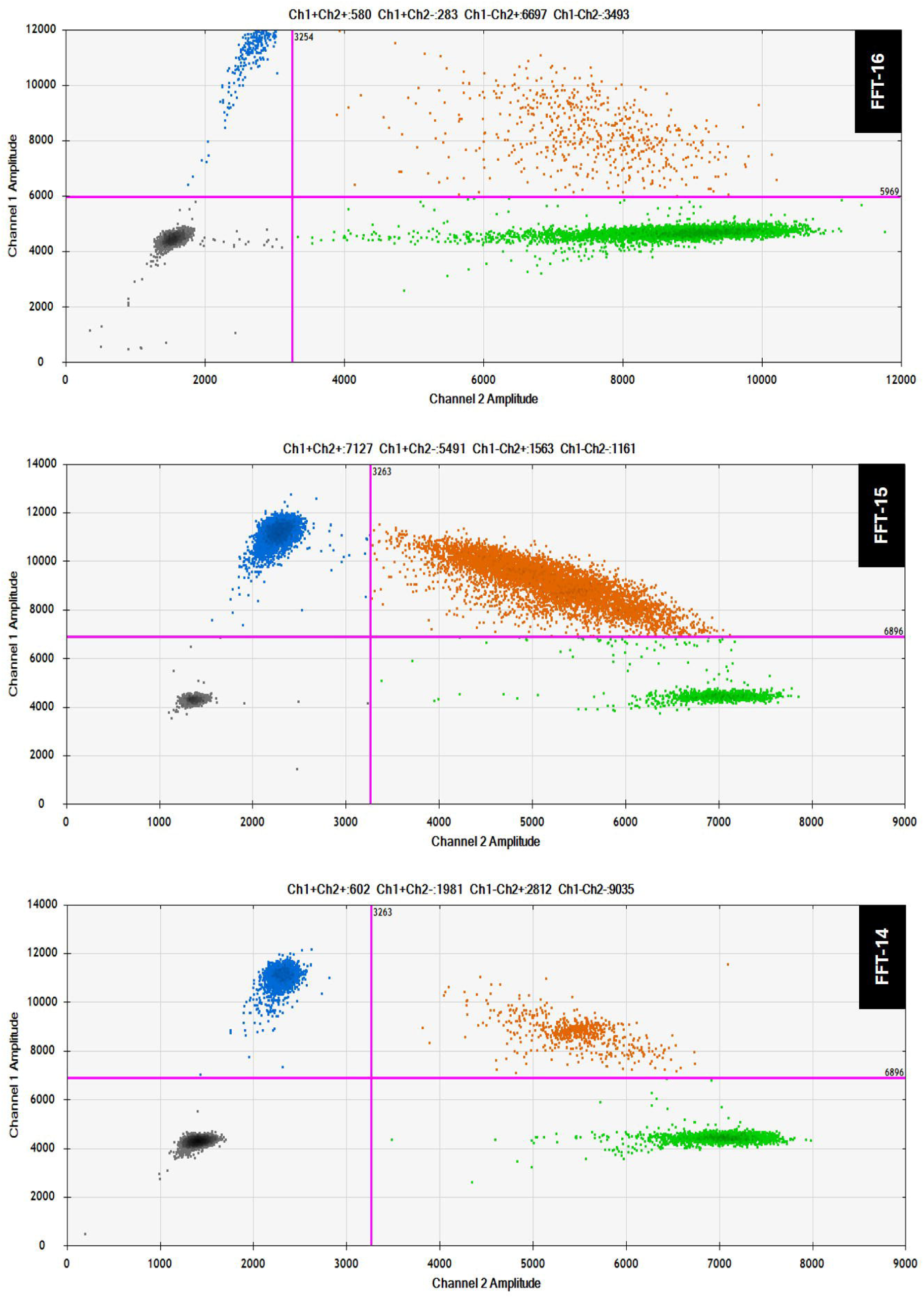

