## Supplementary Information (SI) for "Clamp the LAMP: a photoelectrochemical platform for *KRAS* mutation detection via wild-type blocking"

**Table of Contents**

**Figure S1.** Evaluation of MB scaffolds for the ^1^O_2_-driven PEC platform

**Figure S2.** Validation of cell-line *KRAS* mutation status by Nanopore sequencing

**Figure S3.** Optimization of C-LAMP conditions by gel electrophoresis

**Figure S4.** Clamp incubation temperature tolerance study

**Figure S5.** C-LAMP temperature tolerance study

**Figure S6.** Further optimization of the C-LAMP/^1^O_2_-driven PEC platform

**Figure S7.** *KRAS* copy number analysis of SW620 gDNA by ddPCR

**Figure S8.** Linear regression in mixed WT/mutant DNA and RSD intra-/inter-assay.

**Figure S9.** Validation of healthy controls FFT samples by ddPCR

**Figure S10.** Validation of NSCLC WT FFT samples by ddPCR

**Figure S11.** Validation of NSCLC G12C FFT samples by ddPCR

**Figure S12.** Validation of NSCLC G12V FFT samples by ddPCR

**Table S1.** Genomic characteristics of the cancer cell lines analyzed in this study

**Table S2.** Clinical information of FFT samples analyzed in this study

**Table S3.** List of experimental *KRAS* mutation detection methodologies

**Table S4.** List of gold-standard technologies for *KRAS* mutation detection

**Table S5.** List of oligonucleotides used in this study

**Experimental/methods section**

**References**

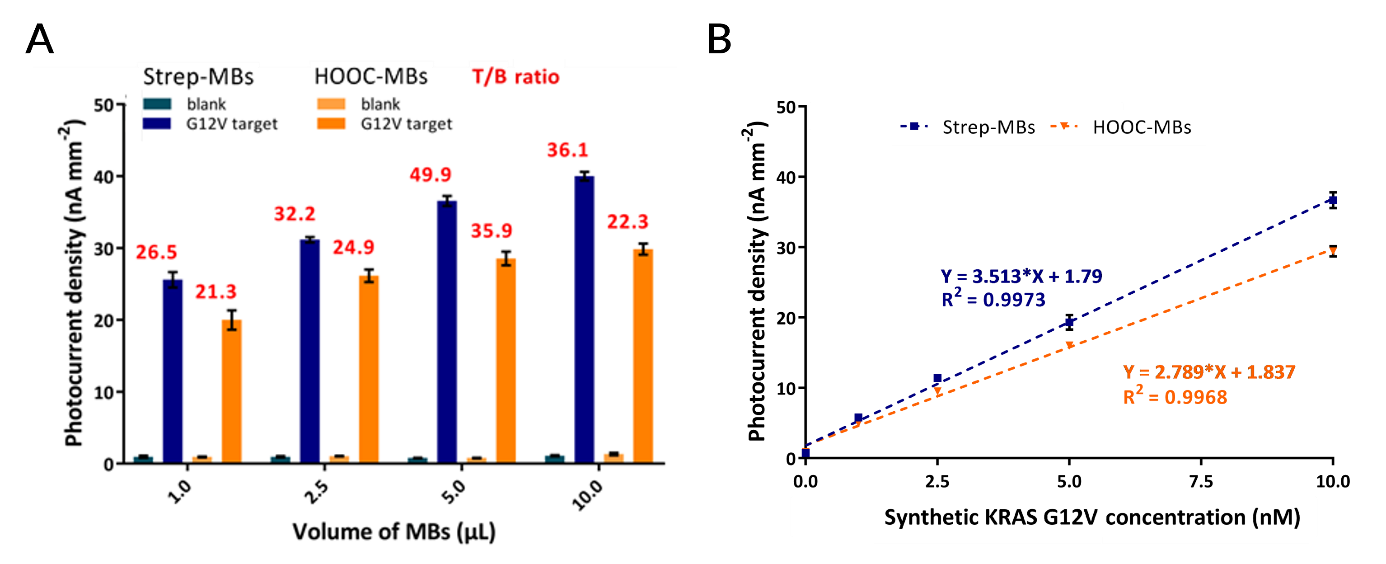

**Figure S1.** Evaluation of MB scaffolds for the ^1^O_2_-driven PEC platform. **(A)** Photocurrent densities measured using 1-10 µL of Strep-MBs bearing b-CPs or HOOC-MBs bearing a-CPs, in the absence (blank) or presence of 10 nM synthetic *KRAS* G12V target. The signal-to-blank ratios are shown in red. **(B)** Calibration plots for *KRAS* G12V detection comparing both MB scaffolds. Error bars represent the standard deviation of the mean (n = 3).

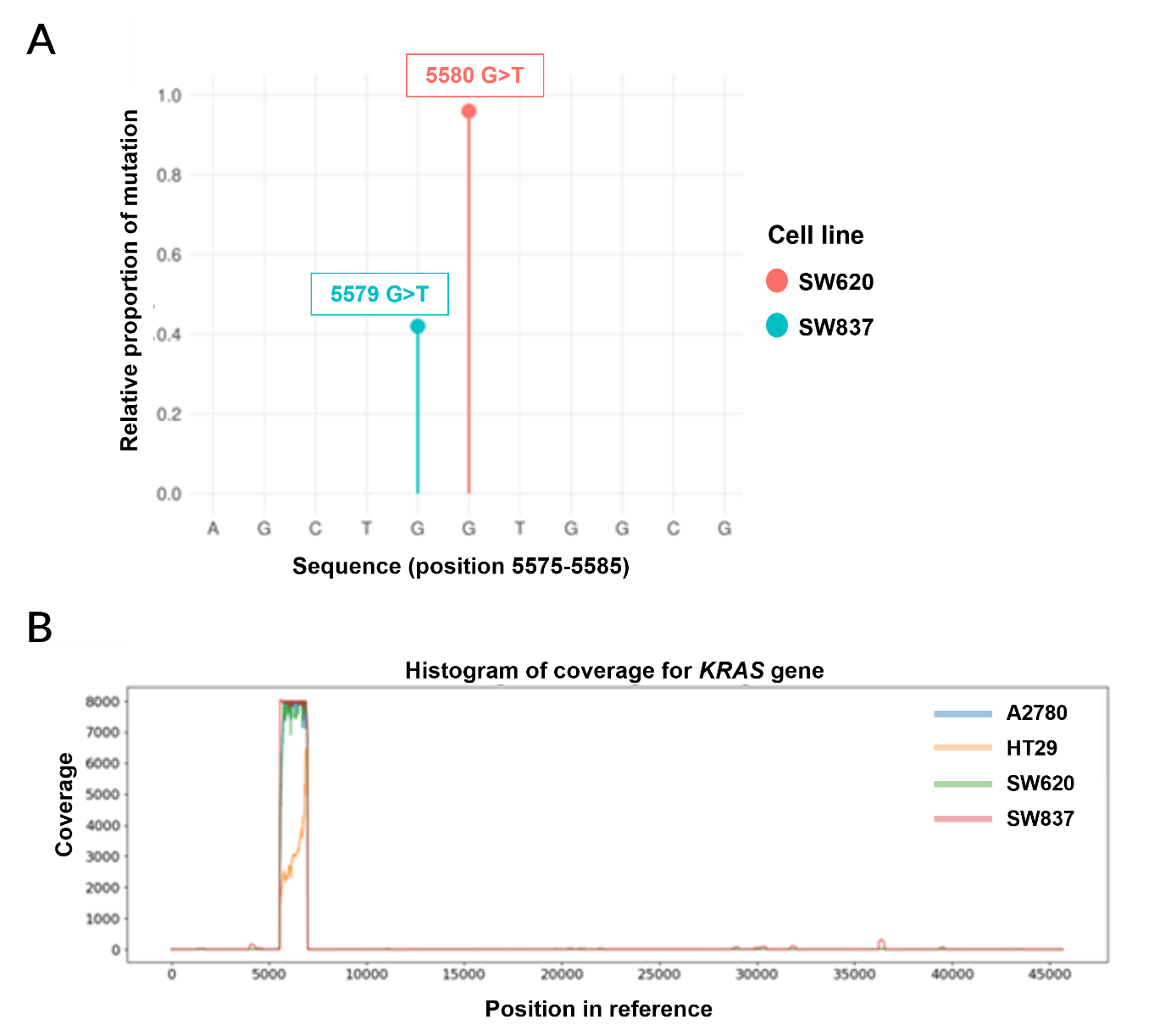

**Figure S2.** Validation of cell-line *KRAS* mutation status by Nanopore sequencing. **(A)** Lollipop plot showing nucleotide substitutions in *KRAS*-mutant cell lines (SW837, G12C/WT; SW620, G12V/G12V). WT cell lines (HT-29 and A2780) showed no mutations at codons 12 or 13 and are therefore absent from the plot. The y-axis indicates the relative proportion of mutated alleles. In nucleotide notation, G12C and G12V correspond to G34T and G35T substitutions, respectively. **(B)** Histogram of sequencing coverage relative to the *KRAS* reference sequence, confirming high read depth within the amplified region between positions 5,500 and 7,000.

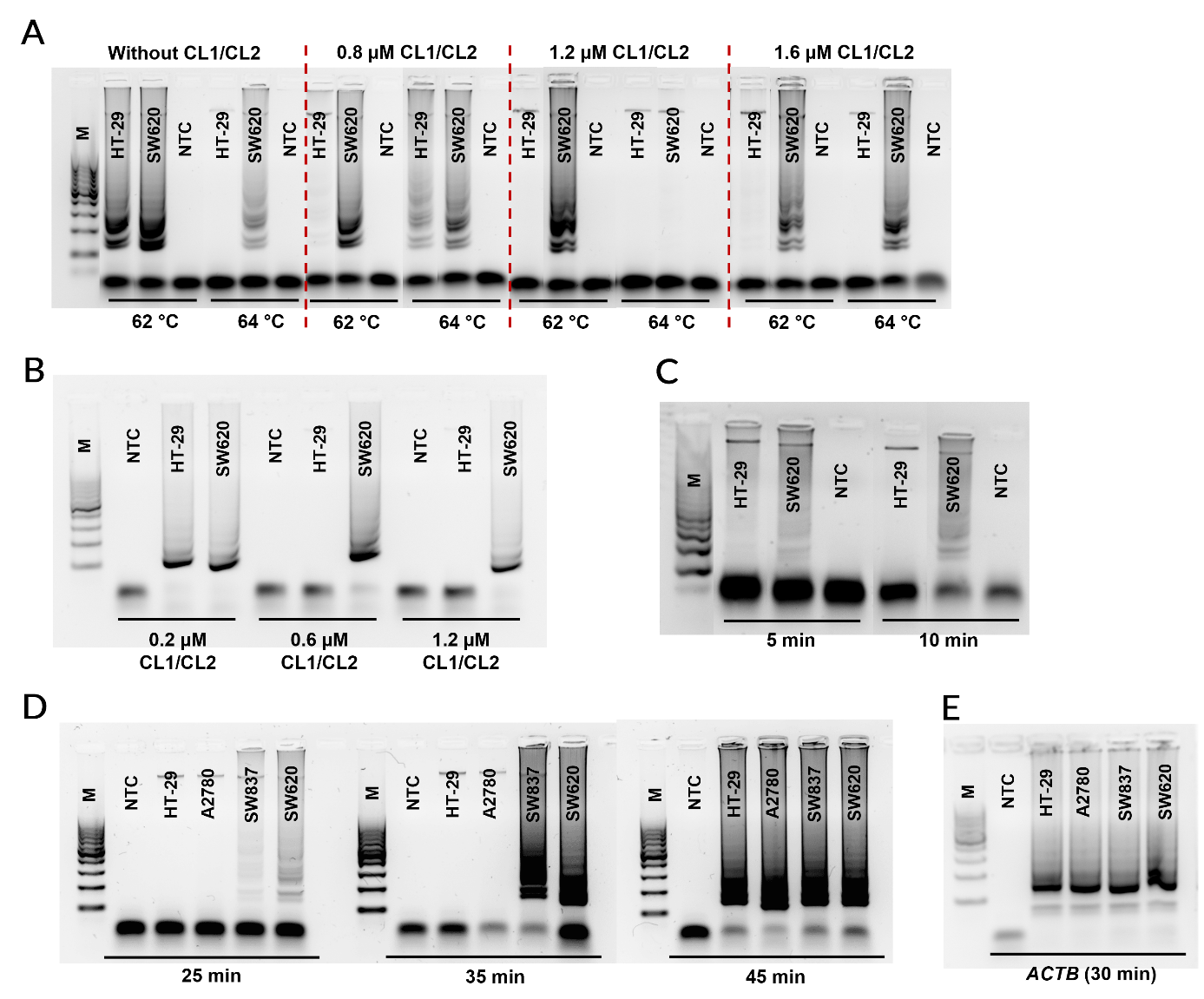

**Figure S3.** Optimization of C-LAMP conditions by gel electrophoresis using NTC, WT (HT29 and A2780) and *KRAS*-mutant (SW837 and SW620) cell lines. **(A-B)** Optimization of clamp probe (CL1/CL2) concentration and reaction temperature identified 0.6 µM at 62 °C as the most effective conditions for WT suppression. **(C)** A 10 min pre-incubation at 40 °C improved clamp hybridization and selectivity. **(D)** Amplification time optimization revealed 35 min as the optimal duration, yielding strong signals for mutant templates while blocking WT amplification. **(E)** *ACTB* amplification, used as an internal control, confirmed DNA quality and integrity across cell lines. M: 100-1,000 bp DNA ladder.

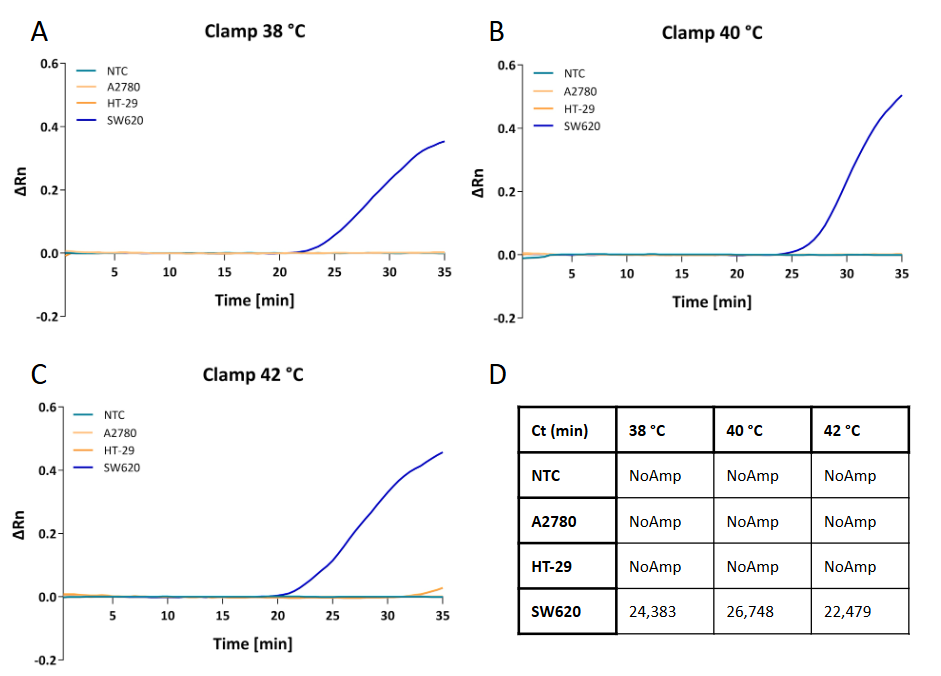

**Figure S4.** Clamp preincubation temperature tolerance study. The DNA with clamp oligonucleotides was incubated at **(A)** 38 °C, **(B)** 40 °C and **(C)** 42 °C for 10 minutes, while all C-LAMP reactions were conducted at 62 °C for 35 minutes. Lines show increase of relative fluorescence of the mutated cell line DNA over time. In some places, lines from NTC, A2780 and HT-29 overlap due to very similar fluorescence signals. **(D)** Table shows final Ct values in minutes. No amplification leakage was observed in wild-type samples across the evaluated ±2 °C preincubation window.

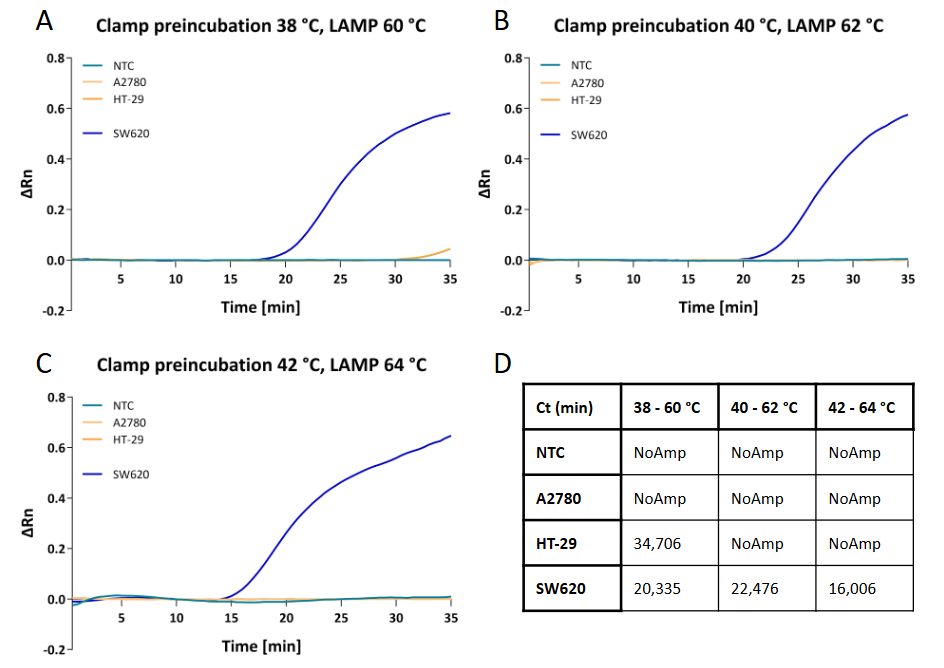

**Figure S5.** C-LAMP temperature tolerance study. The whole C-LAMP protocol temperatures were **(A)** 2 °C lower (38 + 60 °C), **(B)** optimal protocol (40 + 62 °C), and **(C)** 2 °C higher (42 + 64 °C). Lines show increase of relative fluorescence of the mutated cell line DNA over time. In some places, lines from NTC, A2780 and HT-29 overlap due to very similar fluorescence signals. **(D)** Table shows final Ct values in minutes. Wild-type suppression was preserved across the ±2 °C variation range.

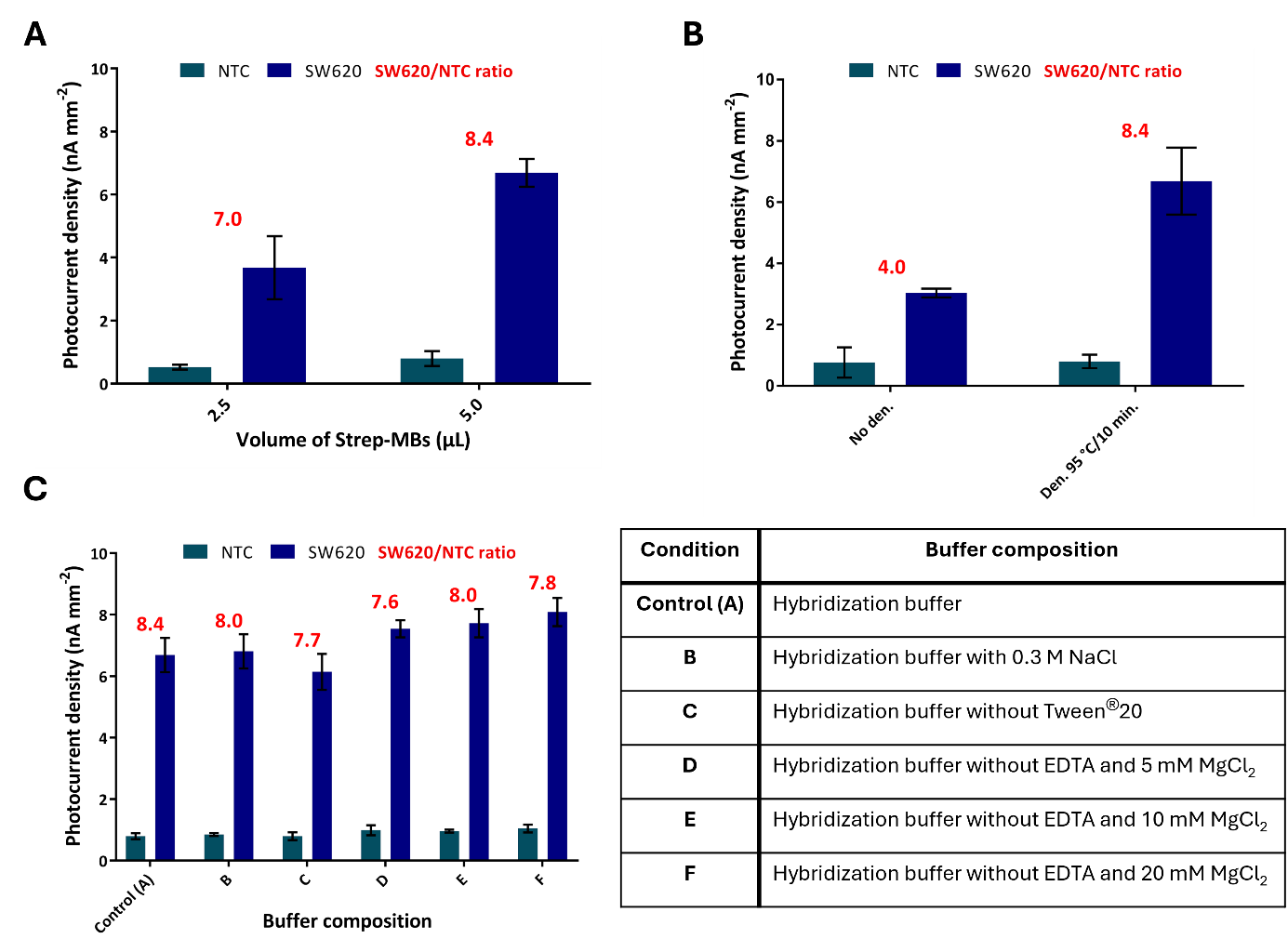

**Figure S6.** Further optimization of the C-LAMP/^1^O_2_-driven PEC platform using gDNA from SW620 (G12V/G12V) and NTC, with b-CPs specific for *KRAS* G12V detection. **(A)** Effect of Strep-MB volume (2.5-5.0 µL). **(B)** Influence of C-LAMP product denaturation on photocurrent response. **(C)** Effect of buffer composition: hybridization buffer (control, bars A), control with 0.3 M NaCl (bars B), control without Tween^®^20 (bars C), control without EDTA and 5 mM (bars D), 10 mM (bars E) or 20 mM (bars F) MgCl_2_. Control condition the in hybridization buffer corresponds to 1 M NaCl. Photocurrent ratios between SW620 and NTC samples are indicated in red. Error bars represent the standard deviation of the mean (n = 3).

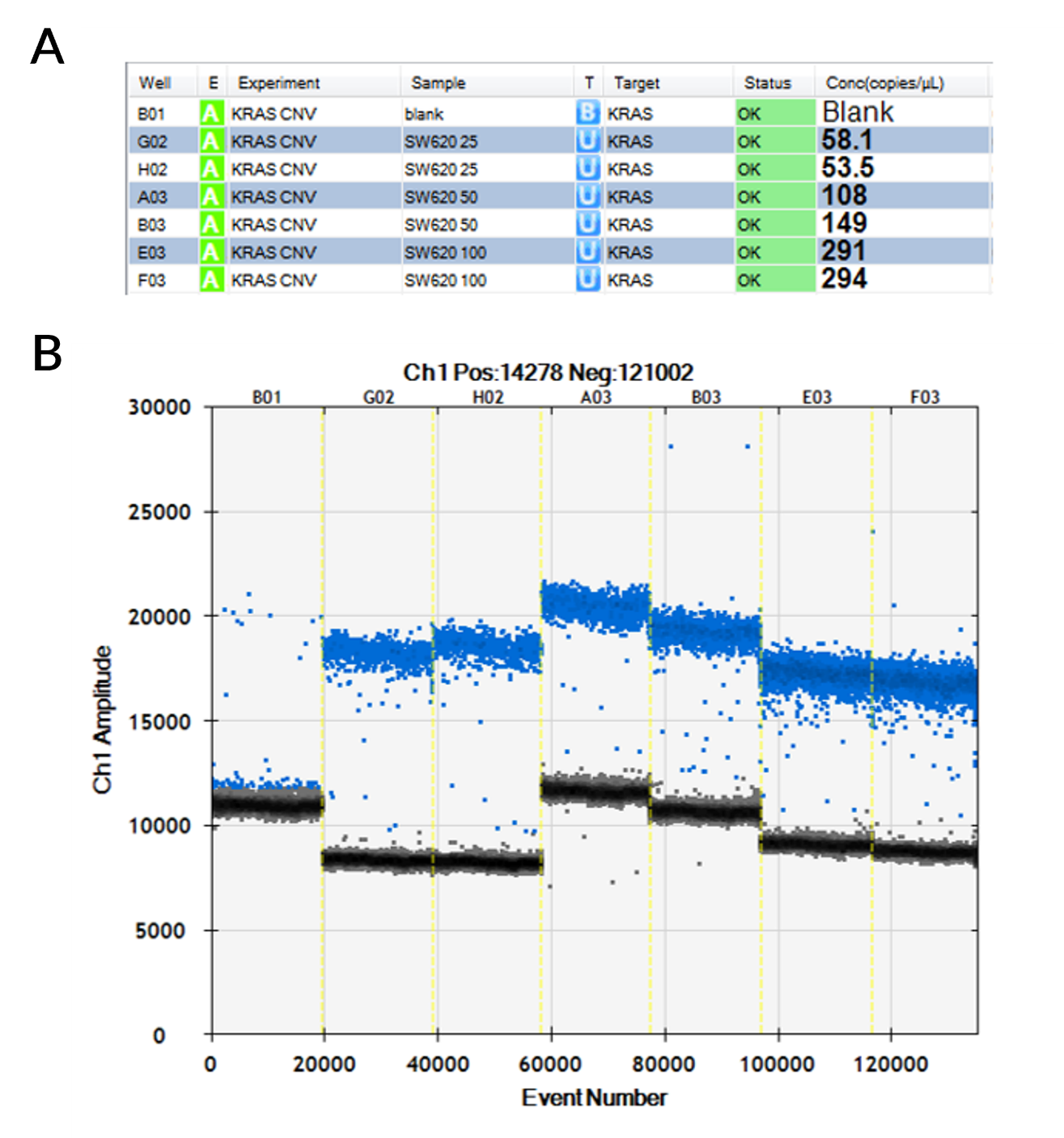

**Figure S7.** *KRAS* copy number analysis of SW620 gDNA by ddPCR. **(A)** Table showing the distribution of samples across plate positions and the calculated *KRAS* copy numbers per microliter of reaction mixture. Three different input amounts of SW620 gDNA were tested to minimize bias from droplet partitioning: 25 ng (SW620 25), 50 ng (SW620 50) and 100 ng (SW620 100). **(B)** One-dimensional amplitude plots showing negative droplets and positive droplets containing *KRAS* gene copies across six replicates. A few positive droplets appeared in the blank sample at a negligible rate. Mean copy number values were used for LOD estimation and molar conversion.

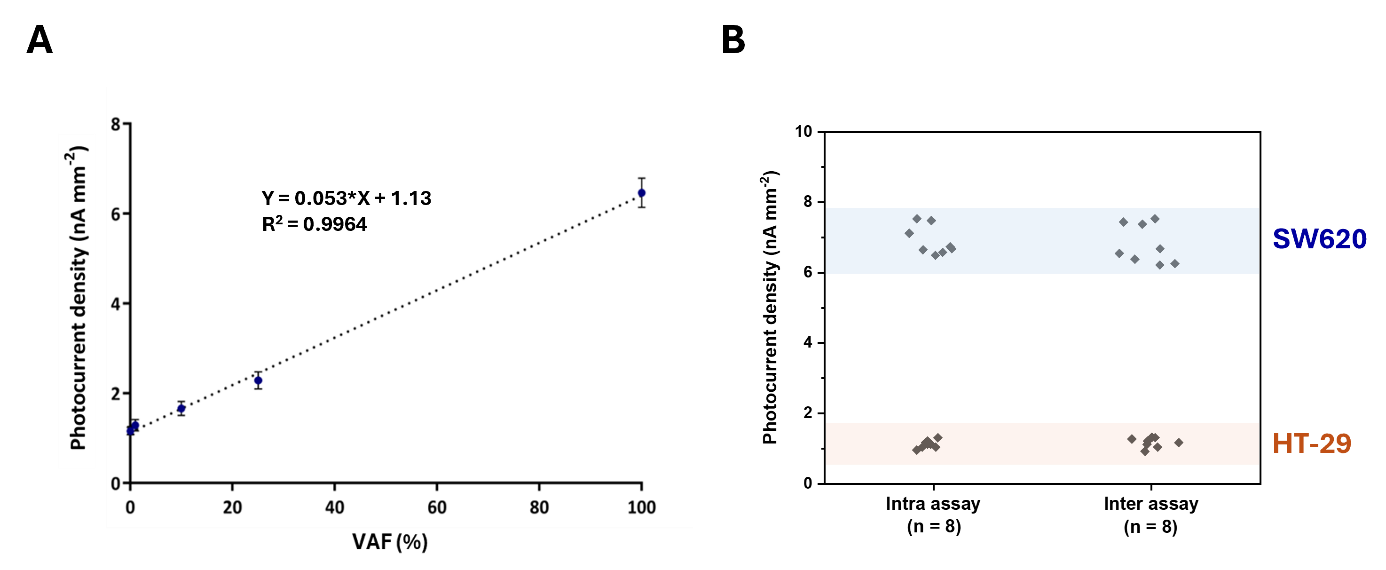

**Figure S8.** Linear regression in mixed WT/mutant DNA and RSD intra-/inter-assay. **(A)** Sensitivity expressed in VAF was evaluated by measurement of SW620 in HT-29 DNA mixture of increasing SW620/HT-29 ratio (VAF) with stable input of 100 ng of total DNA (10 ng µL^-1^ in the C-LAMP reaction). Error bars represent the standard deviation of the mean (n = 3). **(B)** Reproducibility of the complete biosensor fabrication and measurement workflow at clinically relevant DNA concentrations (10 ng µL⁻¹), using independently prepared electrodes and bead suspensions (n = 8).

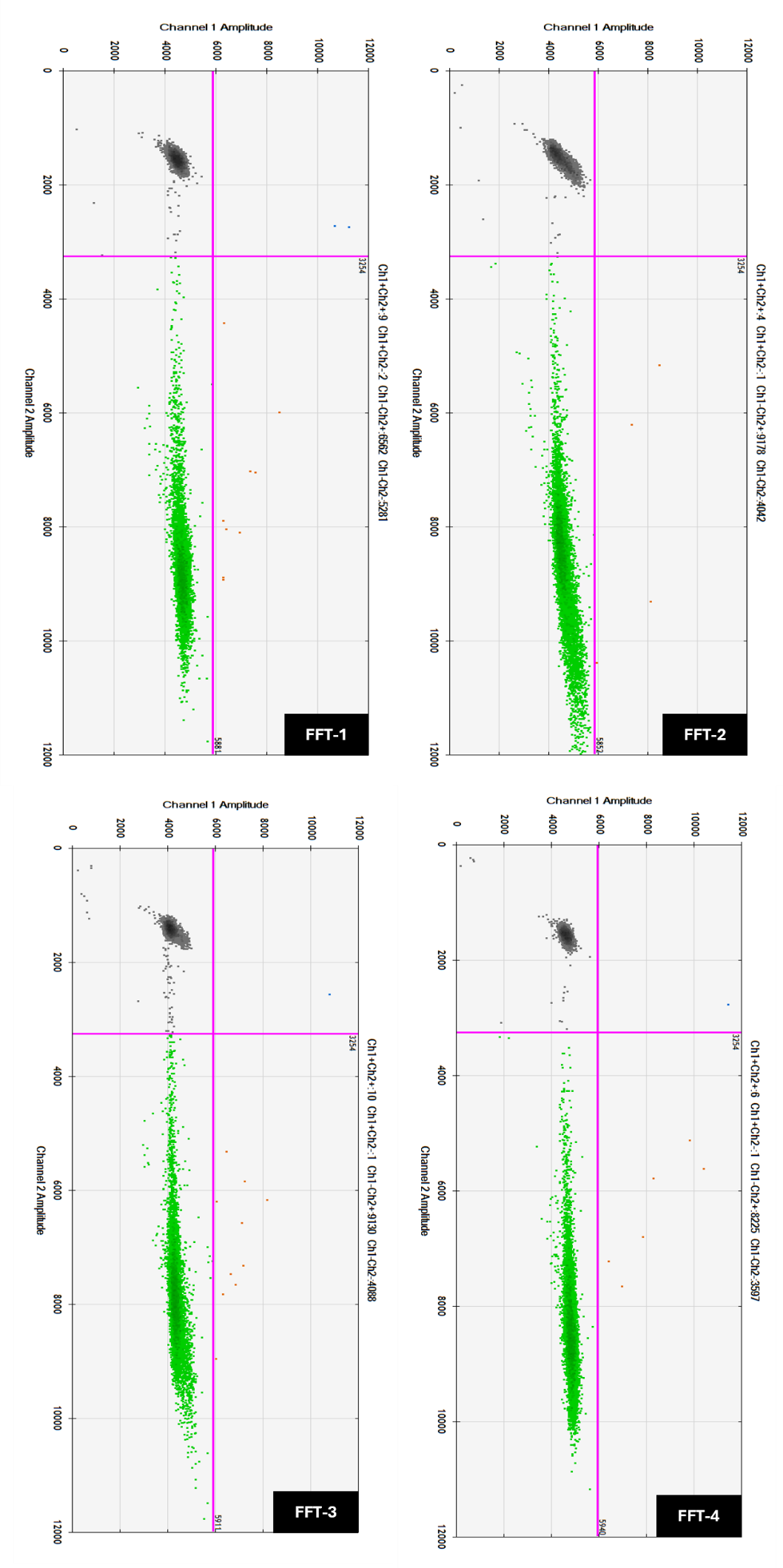

**Figure S9.** Validation of healthy *KRAS* WT FFT samples by ddPCR. Representative two-dimensional amplitude plots showing droplet clustering into negative (black) and WT-positive (green), SNV-positive (blue; G12C or G12V, depending on the sample genotype) and double-positive WT+SNV (orange) populations. FFT-1 to FFT-4 were classified as WT (Table S2), in agreement with parallel NGS analysis performed during standard clinical diagnostics.

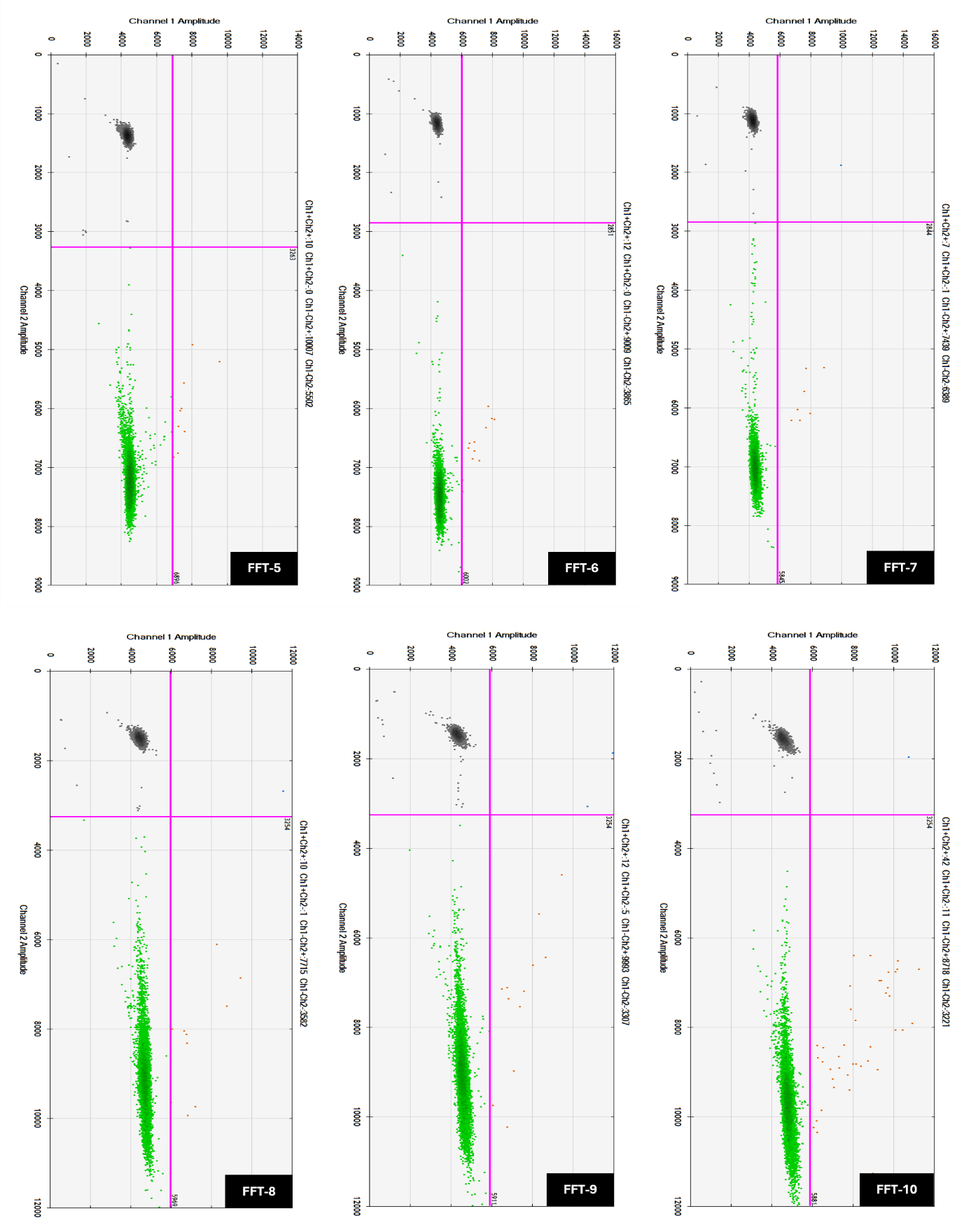

**Figure S10.** Validation of NSCLC *KRAS* WT FFT samples by ddPCR. Representative two-dimensional amplitude plots showing droplet clustering into negative (black) and WT-positive (green), SNV-positive (blue; G12C or G12V, depending on the sample genotype) and double-positive WT+SNV (orange) populations. FFT-5 to FFT-10 were classified as WT (Table S2), in agreement with parallel NGS analysis performed during standard clinical diagnostics.

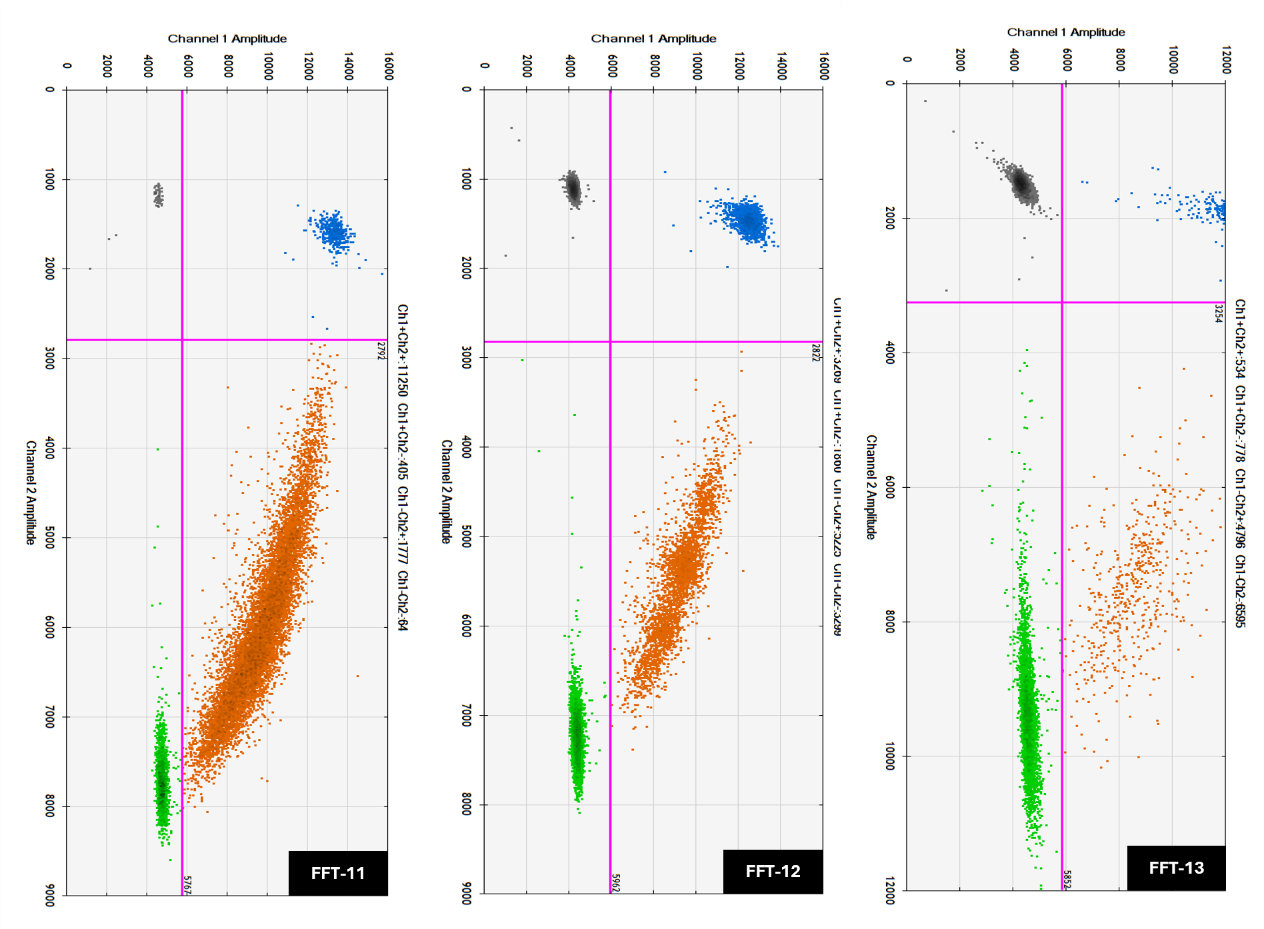

**Figure S11.** Validation of NSCLC *KRAS* G12C FFT samples by ddPCR. Representative two-dimensional amplitude plots showing droplet clustering into negative (black) and WT-positive (green), SNV-positive (blue; G12C or G12V, depending on the sample genotype) and double-positive WT+SNV (orange) populations. FFT-11 to FFT-13 were classified as G12C (Table S2), in agreement with parallel NGS analysis performed during standard clinical diagnostics.

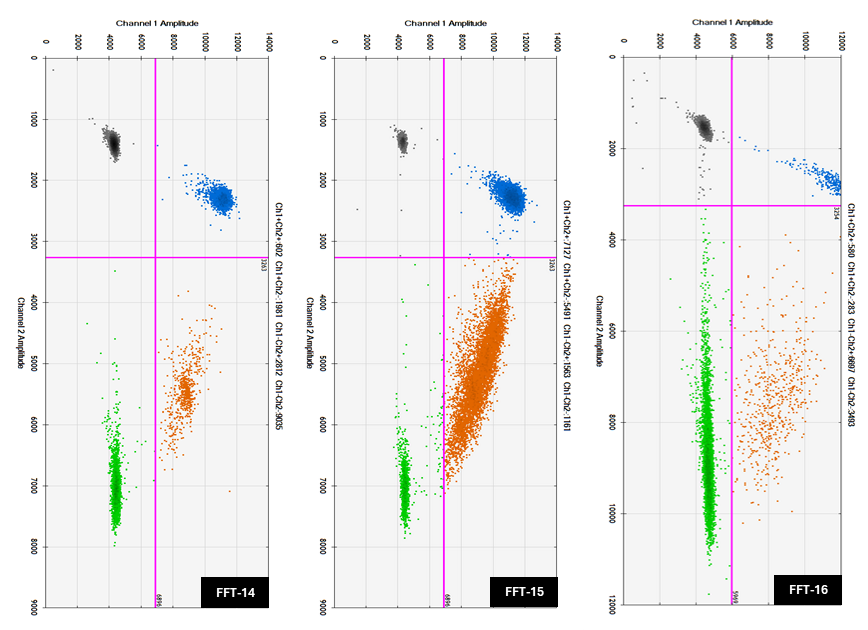

**Figure S12.** Validation of NSCLC *KRAS* G12V FFT samples by ddPCR. Representative two-dimensional amplitude plots showing droplet clustering into negative (black) and WT-positive (green), SNV-positive (blue; G12C or G12V, depending on the sample genotype) and double-positive WT+SNV (orange) populations. FFT-14 to FFT-16 were classified as G12V (Table S2), in agreement with parallel NGS analysis performed during standard clinical diagnostics.

**Table S1.** Genomic characteristics of the cancer cell lines analyzed in this study, summarizing cancer type, codon 12-13 sequences, *KRAS* mutation status and zygosity, as confirmed by Nanopore sequencing. SNVs in codon sequences are underlined. Nucleotide changes are reported following HGVS nomenclature. The selected cell lines were chosen to represent homozygous (WT and mutant) and heterozygous *KRAS* mutation backgrounds.

| **Cell line** | **Cancer type** | **Codon sequence** | **Mutation (nucleotide change)** | ***KRAS* status** | **Zygosity** |
| --- | --- | --- | --- | --- | --- |
| HT-29 | Colorectal | GGTGGC | WT | WT/WT | Homozygous |
| A2780 | Ovary | GGTGGC | WT | WT/WT | Homozygous |
| SW837 | Colorectal | TGTGGC | G12C (c.34G>T) | G12C/WT | Heterozygous |
| SW620 | Colorectal | GTTGGC | G12V (c.35G>T) | G12V/G12V | Homozygous |

**Table S2.** Clinical information of FFT samples analyzed in this study, including sex, age at sample collection, cancer type, stage, *KRAS* mutation, percentage of neoplastic cells and VAF as determined by ddPCR. Nucleotide changes are reported following HGVS nomenclature.

| **Clinical sample** | **Sex** | **Age at sample collection** | **Cancer type** | | **Stage** | ***KRAS* mutation** | | **Neoplastic cells* (%)** | **VAF**  **(%)** |
| --- | --- | --- | --- | --- | --- | --- | --- | --- | --- |
| FFT-1 | M | 79 | | Healthy | - | WT | - | | - |
| FFT-2 | M | 29 | | Healthy | - | WT | - | | - |
| FFT-3 | M | 55 | | Healthy | - | WT | - | | - |
| FFT-4 | F | 71 | | Healthy | - | WT | - | | - |
| FFT-5 | M | 66 | | NSCLC | IA2 | WT | - | | - |
| FFT-6 | F | 60 | | NSCLC | IB | WT | 40-50 | | - |
| FFT-7 | M | 79 | | NSCLC | IB | WT | 80-90 | | - |
| FFT-8 | F | 76 | | NSCLC | IIIA | WT | 60 | | - |
| FFT-9 | M | 78 | | NSCLC | IB | WT | 70-80 | | - |
| FFT-10 | F | 64 | | NSCLC | IA3 | WT | 40-50 | | - |
| FFT-11 | F | 74 | | NSCLC | IB | G12C (c.34G>T) | - | | 37 |
| FFT-12 | M | 72 | | NSCLC | IB | G12C (c.34G>T) | 40-50 | | 32 |
| FFT-13 | M | 67 | | NSCLC | IIB | G12C (c.34G>T) | 10-15 | | 17 |
| FFT-14 | F | 64 | | NSCLC | IA2 | G12V (c.35G>T) | 25-40 | | 42 |
| FFT-15 | F | 65 | | NSCLC | IA2 | G12V (c.35G>T) | 60 | | 33 |
| FFT-16 | M | 75 | | NSCLC | IIIA | G12V (c.35G>T) | 2-5 | | 7 |

*Values estimated by the pathologist to reflect variability in tumor cellularity across representative clinical samples. For FFT-1 and FFT-4, assessment was not possible due to suboptimal slide quality.

**Table S3.** List of experimental *KRAS* mutation detection methodologies.

| **Method** | ***KRAS* mutation** | **LOD** | **Input material** | **Assay time** | **Validation technique** | **Equipment** | **Reference** |
| --- | --- | --- | --- | --- | --- | --- | --- |
| C-LAMP with ^1^O_2_-driven PEC | G12V, G12C | 4.8 % VAF* | 15 samples - FFT DNA | 70 min | ddPCR | Thermoblock, LED source, potentiostat | This work |
| PNA-clamping asymmetric PCR | G12V, G12D | 0.1 % VAF* | 10 samples - FFPE tissues DNA | 2 h | direct Sanger sequencing | qPCR thermal cycler | (Oh et al. 2010) |
| COLD-PCR | G12D, G12V, G12S, G12C, G13D | 2.5 % VAF* | 52 samples - FFPE tissues DNA | 6 h 30 min | Therascreen kit (Qiagen); conv. Sanger sequencing | Thermal cycler, Sanger sequencer | (Carotenuto et al. 2012) |
| Multiplex PCR with bead array | G12A, G12C, G12D, G12R, G12S, G12V, G13D | 1 % VAF* | 140 samples - FFPE tissues DNA | 4 h | ARMS/Scorpion LDT; BigDye terminator sequencing | Thermal cycler, flow cytometer Luminex | (Laosinchai-Wolf et al. 2011) |
| CRISPR/Cas9, RCA, gold nanoparticles detection | G12D | 0.01 % VAF* (G12D), 0.2 fM | Cell lines DNA in FBS | 2 h 30 min | Sanger sequencing, qPCR | Thermoblock | (Zhou et al. 2022) |
| RCA with mutS and ATRP signal amplification | G12D | N/A % VAF*, 3.09 aM | Synth. target spiked in human serum | 3 h | N/A | Thermoblock, EIS device, potentiostat | (Lee et al. 2022) |
| Allele-specific RPA | G12C | 250 copies of gDNA | 10 samples - FFPE tissues DNA | 1 h 30 min | NGS | Thermomixer, microfluidics chip, fluorescence reader | (Martorell et al. 2019) |
| RPA, AS-HCR with smartphone detection | G12C, G13D | 0.7 % VAF* | 36 samples - FFPE tissues DNA | 1 h | NGS | Thermoblock, plastic microarray chips | (Lazaro et al. 2022) |
| RPA, RCA, AI-assisted EC bioassay | G12V | 0.53 % VAF*,  61 pM | 11 samples - FFPE tissues DNA | 3 h | NGS | Multipotentiostat/galvanostat, thermoblock | (Sebuyoya et al. 2025) |
| PNA-LB mediated allele-specific LAMP | G12D | 6.43 ± 3.75 ng of DNA | Cell line DNA | 35 min | Sanger sequencing | Lab-on-a-disc hardware, heating system, visual detection module, | (Mirlohi et al. 2024) |
| PNA-LNA molecular switch + LAMP | G12V, G12D | 25 % VAF*,  1 copy μL^-1^ | 54 samples - 30 FFT DNA, 24 circulating tumor cells DNA | 40 min | Sanger sequencing, HRM analysis | Thermoblock, visual detection, fluorescence reader | (Islam et al. 2025) |
| CRISPR/Cas9 + HCR | G12C, G12D | 0.1 % VAF*,  ~1 fM | spiking - DNA targets in 50% human serum | 4 h 10 min | Sanger sequencing | Thermoblock, fluorescence spectrophotometer, AFM | (Ji et al. 2025) |
| Sticky End-Mediated CRISPR/ Cas12a Coupled RPA | G12C | 0.1 % VAF*,  10 pM | spiking - DNA targets in human serum; cell lines DNA | 1 h 15 min | FastNGS, qPCR, Sanger sequencing | Thermoblock, fluorescence spectrometer | (He et al. 2025) |

* Variant allele frequency

**Table S4.** List of gold-standard technologies for *KRAS* mutation detection.

| **Method** | **LOD** | **Assay time** | **Cost per test**** | **Equipment requirements** |
| --- | --- | --- | --- | --- |
| C-LAMP with ^1^O_2_-driven PEC | 4.8 % VAF* | 70 min | 5 - 6 € | Thermoblock, LED source, potentiostat |
| qPCR | 1 - 5 % VAF* | > 2 h | 3 - 4 € | qPCR thermal cycler, optical readout |
| ddPCR | 1 % VAF* | > 3 h | 6 - 7 € | dPCR thermal cycler, droplet generator/microchamber plates, droplet/plate optical reader |
| NGS (biomarker panel) | 1 - 5 % VAF* | > 1 day | 45 - 90 € | Sequencer including optical detector, thermal cycler |

* Variant allele frequency

** Average price of dedicated reagents in European countries, actual prices may vary depending on the scope of examination and health insurance system. Estimated reagent cost only; instrumentation not included.

**Table S5.** List of oligonucleotides used in this study. Sequences are shown in the 5’→3’ direction, with terminal modifications indicated. LNA modifications are underlined. Ce6, biotin and amino (NH_2_) groups are specified.

| **Name** | **5’ mod.** | **Sequence (5’ → 3’)** | **3’ mod.** |
| --- | --- | --- | --- |
| F3 | – | TTAAAAGGTACTGGTGGAGTA | – |
| B3 | – | ATTGTTGGATCATATTCGTCC | – |
| FIP | – | AGTCATTTTCAGCAGGCCTTATAATTGATAGTGTATTAACCTTATGTGTG | – |
| BIP | – | TAAACTTGTGGTAGTTGGAGCTGGTGGCTGAATTAGCTGTATCGTCAAGG | – |
| BIP.V2 | – | TAAACTTGTGGTAGTTGGAGCTGTGAATTAGCTGTATCGTCAAGG | – |
| BIP.V3 | – | CTTGTGGTAGTTGGAGCTGGTGGCGTGAATTAGCTGTATCGTCAAGG | – |
| CL1 | – | TACGCCACCAGCTCC | -NH_2_ |
| CL2 | – | CCTACGCCACCAGCT | -NH_2_ |
| b-CP (WT) | – | ACGCCACCAGCTCCA | -biotin |
| b-CP (G12V) | – | TACGCCAACAGCTCC | -biotin |
| b-CP (G12C) | – | ACGCCACAAGCTCCA | -biotin |
| a-CP (G12V) | – | TACGCCAACAGCTCC | -NH_2_ |
| G12V target | – | TGGAGCTGTTGGCGTAGGCAAGAGTGCCTTGACGATACAGCT | – |
| DP | Ce6- | AGCTGTATCGTCAAGGCACTCTTGC | – |
| ddPCR Fwd | – | GCCTGCTGAAAATGACTGAATATA | – |
| ddPCR Rev | – | TTAGCTGTATCGTCAAGGCACTC | – |
| SEQ Rev | – | GGAAGCCAAATAAAGTTGTTTC | – |
| *ACTB* F3 | – | GGGCTTCTTGTCCTTTCCTT | – |
| *ACTB* B3 | – | TCAGCAGCACGGGGT | – |
| *ACTB* LF* | – | GCCCACATAGGAATCCTTCT | – |
| *ACTB* LB* | – | ACTGGGACGACATGGAGAAAATCT | – |
| *ACTB* FIP | – | CCTCTCTTGCTCTGGGCCTCGTGTGATGGTGGGCATGGGT | – |
| *ACTB* BIP | – | CATCGAGCACGGCATCGTCACCACACGCAGCTCATTGT | – |

*Additional primers in *ACTB* (LF and LB) were employed following our previous work (Moranova et al. 2024).

**Experimental/methods section**

Additional instrumentation included a horizontal gel electrophoresis unit (Sub-Cell GT Cell, Bio-Rad, USA), a QX200 AutoDG droplet digital PCR system (Bio-Rad, USA) and a T100 thermal cycler (Bio-Rad, USA). Gel images were acquired with a GBOX iChemi system (Syngene, UK). Oligonucleotide concentrations were measured using a NanoPhotometer N60 (Implen, Germany). Magnetic separations were performed with custom-made racks equipped with neodymium magnets (Supermagnete, Belgium). A 914 pH/Conductometer (Metrohm, Switzerland), a digital dry shaking bath and a vortex mixer (ThermoFisher Scientific, USA) were also employed.

Streptavidin-coated magnetic beads (Strep-MBs; Dynabeads® M-280, ф = 2.8 µm, Invitrogen, Thermo Fisher Scientific, USA) and carboxylate-modified magnetic beads (HOOC-MBs; ф = 1 µm, Sera-Mag™, Cytiva, USA) were employed.

All reagents were of analytical grade. Tris-(hydroxymethyl) aminomethane (Tris; VWR, Avantor^®^, USA), magnesium chloride (MgCl_2_), Trizma^®^ hydrochloride (Tris-HCl), potassium phosphate monobasic (KH_2_PO_4_), boric acid (H_3_BO_3_), Tween^®^20 and Titriplex^®^ III disodium salt dihydrate (EDTA) were purchased from Merck (Germany). Hydroquinone (HQ), sodium chloride (NaCl) and agarose were obtained from Thermo Fisher Scientific (USA), while potassium chloride (KCl) was supplied by Union Chimique Belge (Belgium). The Saphir Bst Turbo GreenMaster (STM, Jena Bioscience, Germany) was used as the LAMP Master Mix. For gel electrophoresis, a 100-1,000 bp DNA ladder (Jena Bioscience, Germany), GelRed^®^ nucleic acid stain (Biotium, USA) and loading buffer (TopBio, Czech Republic) were employed. For ddPCR, QX200™ EvaGreen Supermix, Automated Droplet Generation Oil and Droplet Reader Oil (Bio-Rad, USA) were used.

All buffers were prepared with Milli-Q water (18.2 MΩ·cm, Arium^®^ Mini, Sartorius, Germany). The hybridization buffer contained 5 mM Tris-HCl, 0.5 mM EDTA, 1 M NaCl and 0.1% (v/v) Tween^®^20, adjusted to pH 7.5. The measuring solution consisted of 0.1 M KCl and 0.01 M KH_2_PO_4_, pH 7.0. For gel electrophoresis, Tris-borate-EDTA (TBE) buffer was prepared with 100 mM Tris, 100 mM H_3_BO_3_ and 2 mM EDTA, pH 8.0.

All experiments were performed in triplicate (n = 3) and data are expressed as mean ± standard deviation (SD). Error bars in graphs represent the standard deviation of the mean (n = 3).

The photocurrent density (P) was calculated according to Equation 1:

$$P= \frac{I_{signal}-I_{background}}{A}$$

(1)

where *P* is the photocurrent density (nA mm^-2^), *I* the measured current (nA) and *A* the working electrode surface area (mm^2^).

The limit of detection (LOD) was defined as:

$$LOD= \frac{3\sigma}{s}$$

(2)

where *σ* is the SD of blank samples (n = 10) and *s* is the slope of the calibration plot.

The relative standard deviation (RSD) was calculated as:

$$RSD= \frac{SD}{Mean}\times100$$

(3)

DNA copy numbers were converted to attomolar concentrations using Equation 4. For molar conversion, the total C-LAMP reaction volume (10 µL) was considered:

$$C (aM)= \frac{N_{A}\times c \left( {copies \mu L}^{-1} \right)}{V}$$

(4)

where N_A_ is Avogadro’s number (6.022 ∙ 10^23^ mol^-1^), *c* the number of DNA copies per microliter and *V* is the reaction volume (L).

Receiver operating characteristic (ROC) curves were generated by plotting sensitivity against (1 – specificity) across multiple thresholds. The area under the curve (AUC) was calculated to assess diagnostic accuracy, and optimal cut-offs were determined using Youden’s J statistic (J = sensitivity + specificity – 1). Linearity of calibration plots was evaluated by linear regression, and agreement between ^1^O_2_-driven PEC signals and reference assays (ddPCR, Nanopore sequencing) was assessed using Pearson’s correlation coefficient (r).

All statistical analyses were performed using GraphPad Prism 8 (GraphPad Software, USA) and OriginPro 2018 (OriginLab, USA). Figures were prepared using Microsoft Office 365 (Microsoft, USA) and BioRender (BioRender.com).
